# RBM39 shapes innate immunity through transcriptional and splicing control of key factors of the interferon response

**DOI:** 10.1101/2023.10.13.562221

**Authors:** Teng-Feng Li, Paul Rothhaar, Arthur Lang, Oliver Grünvogel, Ombretta Colasanti, Santa Mariela Olivera Ugarte, Jannik Traut, Antonio Piras, Nelson Acosta-Rivero, Vladimir Gonçalves Magalhães, Emely Springer, Andreas Betz, Hao-En Huang, Jeongbin Park, Ruiyue Qiu, Gnimah Eva Gnouamozi, Ann-Kathrin Mehnert, Viet Loan Dao Thi, Stephan Urban, Martina Muckenthaler, Matthias Schlesner, Dirk Wohlleber, Marco Binder, Ralf Bartenschlager, Andreas Pichlmair, Volker Lohmann

**Affiliations:** Department of Infectious Diseases, Molecular Virology, Section Virus-Host-Interactions, Medical Faculty Heidelberg, Heidelberg University, Heidelberg, Germany; Institute of Virology, School of Medicine, Technical University of Munich, Munich, Germany; Department of Infectious Diseases, Molecular Virology, Medical Faculty Heidelberg, Heidelberg University, Heidelberg, Germany; Division of Virus-Associated Carcinogenesis, German Cancer Research Center (DKFZ), Heidelberg, Germany; Institute of Molecular Immunology, University Hospital Klinikum rechts der Isar, Technical University of Munich, Munich, Germany; Bioinformatics and Omics Data Analysis, German Cancer Research Center (DKFZ), Heidelberg, Germany; Heidelberg University, Medical Faculty, Department of Pediatric Oncology, Hematology, Immunology and Pneumology, Heidelberg, Germany; Department of Infectious Diseases, Virology, Medical Faculty Heidelberg, Heidelberg University, Heidelberg, Germany; German Center for Infection Research (DZIF), Heidelberg Partner Site, Heidelberg, Germany; Biomedical Informatics, Data Mining and Data Analytics, Faculty of Applied Computer Science and Medical Faculty, University of Augsburg, Augsburg, Germany; German Center for Infection Research (DZIF), Munich Partner Site, Munich, Germany

**Author notes:** Corresponding author’s. **Author Contributions:** Conceptualization: VL, TL, OG, JT Methodology: VL, OG, OC, NAR, GEG, AKM Investigation: TL, AL, JT, SMOU, PR, VGM, APir, ES, RQ, AB, HEH, JP, Visualization: TL, PR, OG, JT, Funding acquisition: VL Project administration: VL Supervision: VL, AP, RB, MB, DW, MS, MM, SU, VLDT Writing – original draft: TL, OC, VL Writing – review & editing: all authors. **Competing Interest Statement:** Authors declare that they have no competing interests.

**Keywords:** RBM39, IRF3, IFNs, splicing

## Abstract

RNA-binding motif protein 39 (RBM39) is an RNA-binding protein involved in tumorigenesis, cell metabolism, and development. Here, we performed a genome-wide CRISPR/Cas9 screen in two liver-derived cell lines and identified RBM39 as a regulator of cell intrinsic innate immune responses. The knockdown of *RBM39* or the treatment with Indisulam, an aryl sulfonamide drug targeting RBM39 for proteasomal degradation, strongly reduced the induction of interferon-stimulated genes (ISGs) in response to double-stranded RNA (dsRNA) or viral infections upon sensing by toll-like receptor 3 (TLR3) or cytosolic RIG-I-like receptors. RNA sequencing (seq) and mass spectrometry identified that transcription and/or splicing of the key pathway components IRF3, RIG-I, and MDA5 were affected by RBM39 depletion. *RBM39* knockdown further restrained type I and type III IFN pathways, by reducing expression of the type I IFN receptor subunit interferon alpha and beta receptor subunit 2 (IFNAR2), type III IFN receptor subunit interleukin 10 receptor subunit beta (IL-10RB) and transcription factor signal transducer and activator of transcription (STAT) 1 and 2. RBM39 overall orchestrates innate immunity by regulating basal expression of key factors of the interferon response via transcription and/or alternative splicing.

**Significance:** The function of RBM39 in tumorigenesis has been investigated intensively in the last decade, but its immunological role is still largely unknown. In our study, we identified RBM39 as a regulatory factor of cell intrinsic signaling via a CRISPR/Cas9 screen. Depletion of RBM39 impairs TLR3, RIG-I/MDA5, and IFN pathways, and thus attenuates innate immune responses. Our omics analysis revealed that RBM39 governs the basal expression of several key factors within these pathways, such as RNA sensors RIG-I and MDA5, type I/III receptors, transcription factors IRF3, STAT1 and STAT2, via its transcriptional and splicing function. Therefore, RBM39 might be a therapeutic target to modulate innate immunity, e.g. in the context of autoimmune disorders.

## Introduction

The cell intrinsic innate immune system belongs to the first line of host defense mechanisms protecting against invading pathogens. When infected, a limited number of host Pattern Recognition Receptors (PRRs) plays a vital role in recognizing several Pathogen-associated molecular patterns (PAMPs), which activate innate immune responses through complex signaling cascades to eliminate infections. Viral components, including viral DNA (*1, 2*), single-stranded RNA (ssRNA) (*3, 4*), double-stranded RNA (dsRNA) (*5, 6*) and virus-encoded proteins (*7*), are sensed by distinct PRRs. The recognition triggers the cell intrinsic innate immune response in infected cells. For instance, dsRNA, a replication intermediate of many RNA viruses, is sensed by cytosolic retinoic acid-inducible gene (RIG)-I-like receptors (RLRs, RIG-I and MDA5) (*5, 6, 8*) and endosomal Toll-like receptor 3 (TLR3) (*9*). These PRRs then recruit the adaptor proteins mitochondrial antiviral signaling protein (MAVS) and TIR domain containing adaptor molecule 1 (TRIF), respectively, activating a chain of kinases, E3 ubiquitin ligases and E2 ubiquitin-conjugating enzymes. This leads to nuclear factor kappa B subunit 1 (NF-κB) pathway activation and to production of pro-inflammatory cytokines or phosphorylating IRF3, leading to its dimerization and translocation from cytoplasm into nucleus. Subsequently, IRF3 binds to responsive promoters and induces transcription of type I/type III interferons (IFNs) and interferon stimulated genes (ISGs). IFNs act in an auto- and paracrine manner, binding to their receptors, Interferon alpha and beta receptor (IFNAR) and Interferon lambda receptor (IFNLR), on the cell surface, respectively, but signaling via a common cascade upon phosphorylation of signal transducer and activator of transcription 1 and 2 (STAT1 and STAT2). Eventually, STATs form hetero- or homodimers and translocate into nucleus to accelerate or initiate ISG expression in infected and neighboring cells, respectively.

RNA-binding motif protein 39 (RBM39) (also known as HCC1.4 or CAPERα) was initially identified as an autoantigen in a hepatocellular carcinoma patient (*10*) and has been found to be upregulated in multiple cancer cells (*11–13*). It shares high similarity with the splicing factor U2AF 65 kDa subunit (U2AF65), and contains an N-terminal serine/arginine (SR)-rich domain followed by two RNA-recognition motifs (RRM) and a third C-terminal noncanonical RRM belonging to the U2AF homology motif (UHM) family, which mediates the interaction with UHM-ligand motif (ULM) of U2AF65 (*14*). Previous studies have demonstrated that RBM39 functions as a coactivator of activating protein-1 (AP-1), Estrogen Receptor 1(ERα) and Estrogen Receptor 2 (ERβ) (*15–17*) and that it is involved in multiple biological processes, including cell cycle control (*18, 19*), cell metabolism (*19*), and tumorigenesis (*20, 21*), possibly through its splicing activity. In recent years, aryl sulfonamide drugs, such as Indisulam and E7820, were found to degrade RBM39 in a DDB1 and CUL4 associated factor 15 (DCAF15)-dependent manner and thus have been widely investigated in anticancer research (*22–24*). While most studies on RBM39 so far focused on cancer cells, there is limited knowledge of its function in immunity, with one *in vivo* study suggesting that Indisulam-mediated modulation of RBM39 enhances anti-tumor immunity (*25*).

In our study, RBM39 emerged as a prominent candidate through a CRISPR/Cas9-mediated screening in two liver-based cell lines aimed at identifying novel factors associated with the TLR3 response in hepatocytes. Our study reveals that RBM39 exerts a broader influence on innate immunity by modulating the transcription and splicing processes involved in the basal expression of IRF3 and other crucial factors in the IFN signaling pathway. Based on the findings, it is plausible to consider that drugs targeting RBM39, such as Indisulam, may function as immunosuppressants in individuals undergoing treatment.

## Results

### A genome-wide CRISPR/Cas9 Screen identifies novel factors contributing to TLR3 signaling

The initial aim of this study was the identification of novel factors contributing to the regulation of cell intrinsic innate immune responses, in particular focusing on TLR3-mediated response in hepatocytes.

To establish a positive selection system, we developed a death reporter system expressing the suicide gene truncated BH3 interacting domain death agonist (tBID) under the transcriptional control of the interferon induced protein with tetratricopeptide repeats 1 (IFIT1) promoter (Fig. 1A). Upon activation of the TLR3 pathway, tBID expression is induced, resulting in apoptosis (*26*). For the screening we chose PH5CH (*27*), a non-neoplastic, immortalized hepatocyte cell line expressing all PRRs responsive to dsRNA, and Huh7-Lunet-TLR3 (*28*), a hepatocellular carcinoma (HCC)-derived cell line requiring ectopic expression of PRRs for efficient response to dsRNA. The tBID reporter was transduced into both cell lines for stable expression. To evaluate the approach, TLR3-specific stimulation was achieved by the addition of the synthetic dsRNA analog polyinosinic:polycytidylic acid (poly(I:C)) to the cell culture supernatant, leading to efficient tBID-associated, TLR3-dependent cell death only upon activation of the *IFIT1* promoter in both cell lines, in contrast to the empty vector (Fig. S1A). Subsequently, we transduced the IFIT1-tBID cell lines with lentiviral vectors encoding Cas9 along with the GeCKOv2.0 human genome-wide single guide RNA (sgRNA) library, that includes a total of 122,411 unique sgRNAs targeting 19,050 genes and 1,864 miRNAs (Fig. 1B). After selection with puromycin, we added 50 µg/ml poly(I:C) to the medium to activate the TLR3 pathway, or PBS as a control. As a consequence, cells with an intact TLR3 pathway were killed due to the induction of tBID expression, whereas cells expressing guide RNAs targeting genes involved in TLR3 signaling survived, resulting in the enrichment of such sgRNAs. We then analyzed the sgRNA population of stimulated and non-stimulated cells at 72 hours post-stimulation by next-generation sequencing (NGS), to identify target genes interfering with the TLR3 pathway.

**Fig. 1.**
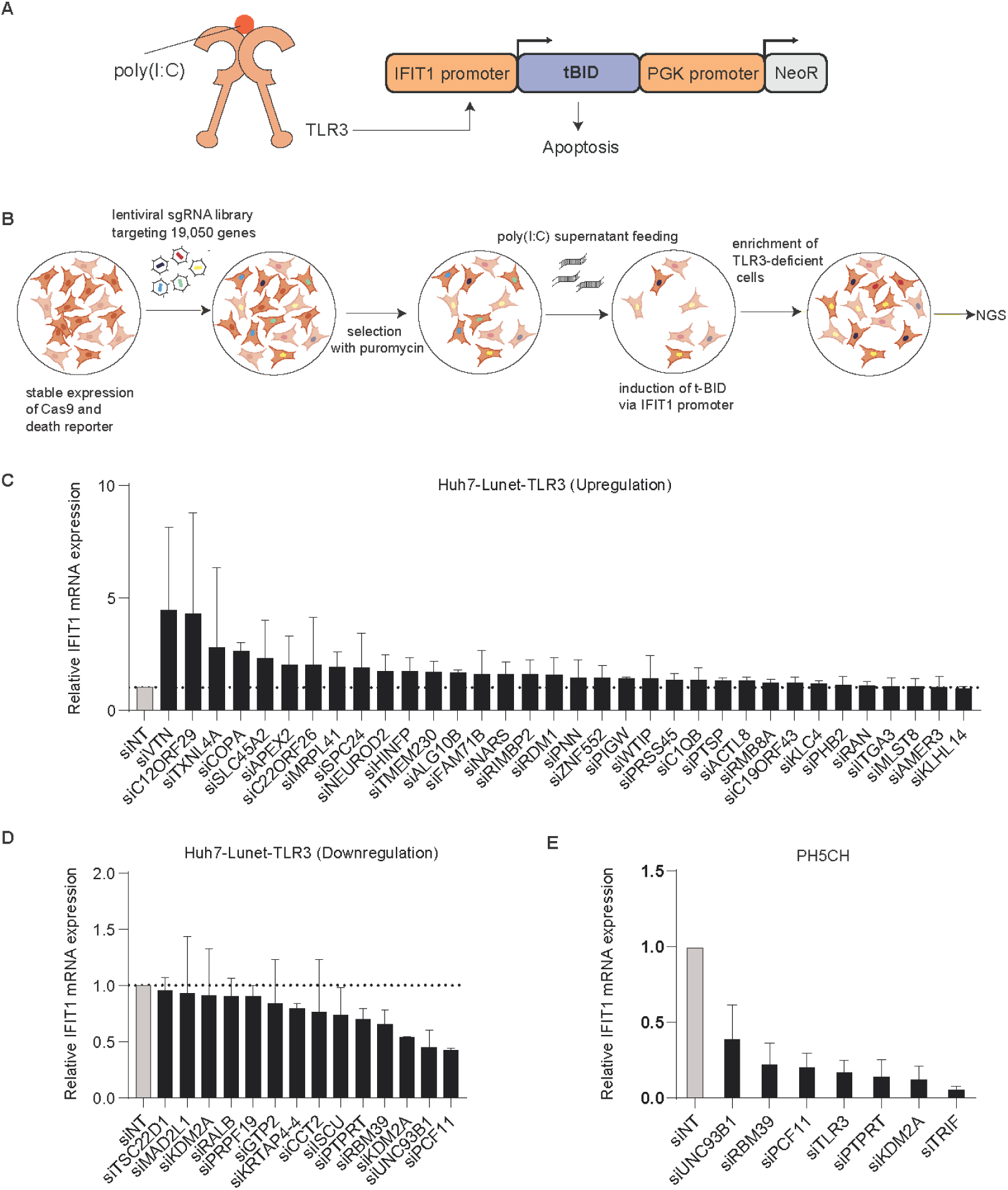
A genome-wide CRISPR/Cas9 screen identifies potential TLR3 related factors. (**A**) Schematic of the truncated BID (tBID) death reporter system. tBID expression and neomycin resistance (NeoR) are under control of the *IFIT1* promoter and the constitutive PGK promoter, respectively. Upon the stimulation by poly(I:C), the *IFIT1* promoter is activated via TLR3 signaling, inducing tBID and resulting in apoptosis. (**B**) Workflow of the CRISPR/Cas9 screen. PH5CH and Huh7 Lunet-TLR3 stably expressing the death reporter and Cas9 were transduced with the genome-wide lentiviral CRISPR sgRNA library and selected with puromycin for stable expression. TLR3 stimulation with poly(I:C) induced expression of tBID and thus apoptosis in case of intact TLR3 signaling. Then, surviving TLR3-deficient cells were enriched and collected for Next Generation Sequencing (NGS). (**C** to **E**) 50 candidate genes were selected and subjected to siRNA silencing for 48 h; then the cells were stimulated by 50 μg/ml poly(I:C) supernatant feeding for 8 h. *IFIT1* mRNA was measured by RT-qPCR. Knockdown of the candidates up-regulated (**C**) or down-regulated (**D**) *IFIT1* mRNA expression in Huh7 Lunet-TLR3 cells. Knockdown of *RBM39, PCF11, PTPRT* and *KDM2A* in PH5CH cells reduced *IFIT1* mRNA expression in PH5CH cells (**E**). siUNC93B1, siTLR3 and siTRIF were used as positive controls. Relative expression to siNT-treated samples was normalized on GAPDH. Data are from three biological replicates (n = 2), error bars indicate standard deviation (SD).

Candidate genes were determined by being enriched beyond a cell-line-specific threshold calibrated to yield about 2,000 overrepresented sgRNAs for each cell line. Due to the expected background of the screen, we needed to narrow down the enriched genes to obtain primary screen hits. We filtered primary candidate genes using a thresholding strategy that was calibrated to include three positive controls (TIR-domain-containing adapter-inducing interferon-β (TRIF), TLR3 and unc-93 homolog B1 (UNC93B1)) for each cell line (*29–31*). A gene was considered as a primary hit in case 5 out of 6 different sgRNAs found enriched in the poly(I:C) treated samples of at least one replicate and the presence of at least one sgRNA targeting a gene in 2 out of 3 replicates. After such filtering, 50 protein-coding genes were chosen for functional validation, whereas 5 microRNAs were excluded despite matching our criteria (Table S1). The protein-coding genes were validated by siRNA knockdown in Huh7-Lunet-TLR3 cells (Fig. 1C and D). However, knockdown of most candidates resulted in unimpaired or even upregulated TLR3 responses (Fig. 1C), likely due to a high background of surviving cells even in absence of sgRNAs (Fig. S1A), resulting in non-specific enrichment. Only the knockdown of a few genes reduced the TLR3 response to levels comparable to the positive control (siUNC93B1). The four candidates with strongest phenotype, RBM39, lysine demethylase 2A (KDM2A), protein tyrosine phosphatase receptor type T (PTPRT), and cleavage and polyadenylation factor II subunit (PCF11) were chosen for further validation in PH5CH cells.

Indeed, knockdown of these factors strongly attenuated innate immune responses in PH5CH cells as well (Fig. 1E). However, PCF11 was not followed up due to its general role in termination of transcription (*32*). *PTPRT* mRNA turned out not to be detectable in Huh7 or PH5CH cells, and the overexpression of siKDM2A-resistant mutant in PH5CH cells failed to rescue the KDM2A phenotype (Fig. S1B), arguing for off-target effects of these siRNAs, therefore RBM39 was chosen for further studies.

### RBM39 knockdown reduces IRF3-but not NF-κB-induced genes upon TLR3 activation

Since PH5CH was considered the most authentic among the available hepatocyte models due to its non-neoplastic nature, we decided to continue here with an in-depth analysis of the specific role of RBM39 on TLR3-IRF3 signaling (Fig. 2A).

**Fig. 2.**
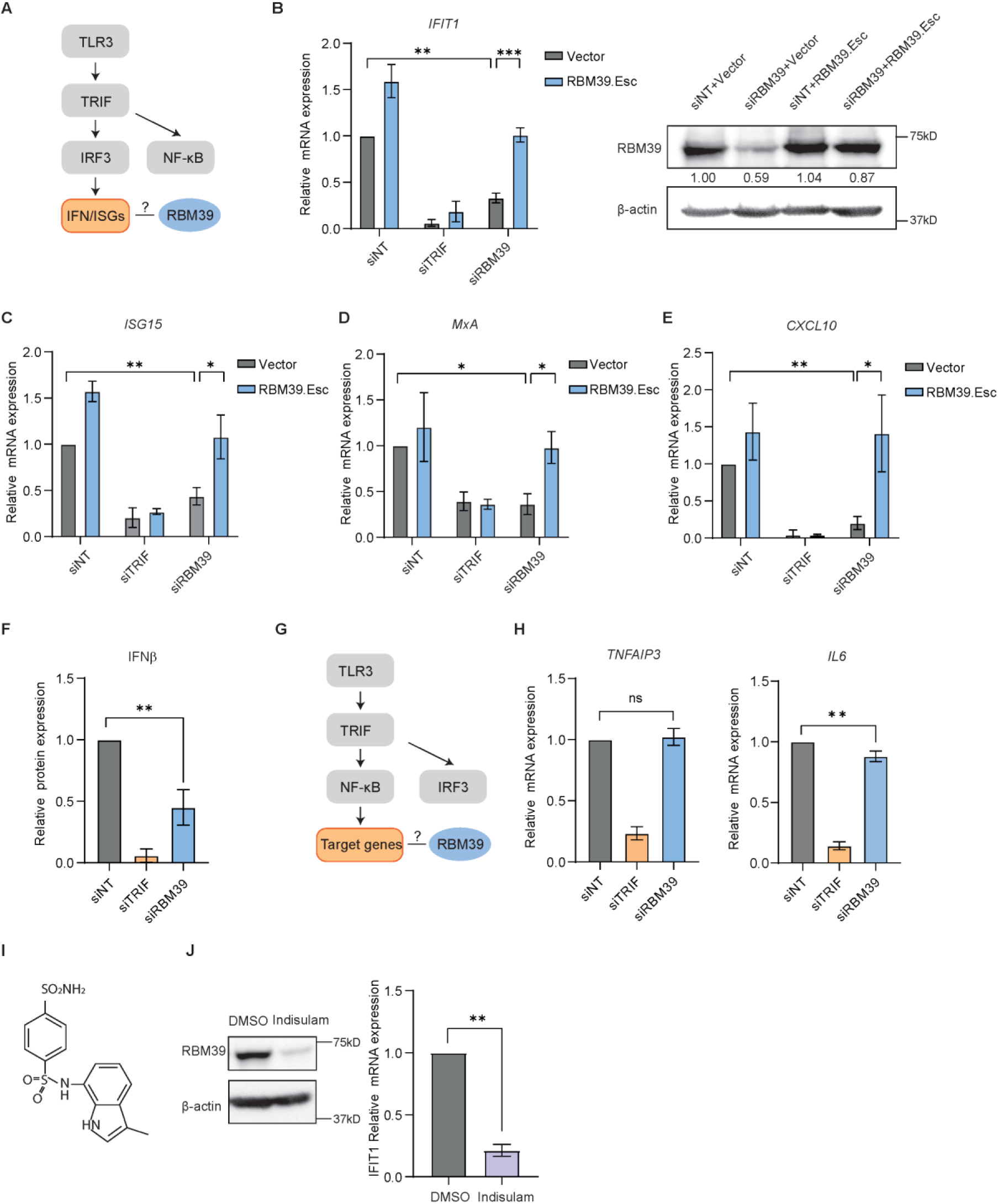
RBM39 is crucial for the TLR3 pathway. (**A**) Simplified schematic of the TLR3-IRF3 pathway. (**B** to **E**) RBM39 rescue experiment. PH5CH cells expressing RBM39.Esc or empty vector were transfected with siRBM39 or siNT/siTRIF as controls for 48 h, and then supernatant-fed with 50 µg/ml poly (I:C) for 6 h. *IFIT1* (**B**), *ISG15* (**C**), *MxA* (**D**) and *CXCL10* (**E**) mRNA was measured by RT-qPCR, RBM39 protein expression was measured by immunoblotting (**B**, right panel). (**F** to **H**) PH5CH cells were transfected with siRNA for 48h and then fed with 50 μg/ml poly(I:C) in the supernatant for 6 h. Secreted IFNβ protein (**F**) was measured by ELISA. Simplified schematic of the TLR3-NF-kB pathway (**G**). TNFAIP3 (**H**, left) and IL6 (**H**, right) mRNA was quantified by RT-qPCR. (**I**) The chemical structure of Indisulam. (**J**) PH5CH cells were treated with 1 µM Indisulam or DMSO as control for 48 h, and further stimulated with 50 µg/ml poly (I:C) in the supernatant for 6 h. RBM39 degradation was measured by western blot (left). *IFIT1* mRNA expression was measured by RT-qPCR (right). Relative expression to siNT control was shown, mRNA fold change was normalized on *GAPDH*. Data are obtained from three biological replicates (n = 3), error bars refer to SD. Statistical significance was assessed through Welsch’s unpaired *t*-test.

First of all, an RBM39 siRNA-resistant mutant (RBM39.Esc) was able to rescue the *IFIT1* mRNA expression upon siRNA knockdown in both PH5CH (Fig. 2B) and Huh7-Lunet-TLR3 cells (Fig. S2A), demonstrating that the phenotype indeed was mediated by RBM39 knockdown. To rule out that RBM39 might have an exclusive impact on *IFIT1* expression, which so far served as the sole readout in our validation, we further analyzed modulation of other ISGs upon *RBM39* knockdown and TLR3 stimulation, including Interferon-Stimulated Protein, 15 KDa (ISG15), MX dynamin like GTPase 1 (MxA) and C-X-C motif chemokine ligand 10 (CXCL10) in PH5CH cells. All tested ISGs showed reduction in *RBM39*-silenced cells compared to the control, and RBM39.Esc overexpression again rescued the knockdown effect (Fig 2 C-E), excluding the off-target effect of siRBM39 and suggesting that RBM39 has a broad impact on various ISGs. To confirm this on protein level, we additionally detected IFN-β secretion via ELISA upon *RBM39* knockdown and observed a significant decline here as well (Fig. 2F).

Considering that the activation of PRRs not only leads to production of various ISGs, but also triggers NF-κB signaling, resulting in the production of cytokines and chemokines, we investigated a potential involvement of RBM39 in the NF-κB pathway (Fig. 2G). However, the expression of two NF-κB-induced genes, tumor necrosis factor alpha induced protein 3 (TNFAIP3) and interleukin 6 (IL6) was not or only mildly affected in PH5CH cells upon poly(I:C) activation and *RBM39* knockdown (Fig. 2H), indicating a minor effect of RBM39 on the NF-κB pathway in this experimental setting.

Indisulam is a sulfonamide drug mediating the interaction between the E3 ligase DCAF15 and RBM39, leading to the ubiquitination and proteasomal degradation of RBM39 (Fig. 2I) (*22, 33, 34*). Indeed, Indisulam dramatically reduced RBM39 expression in PH5CH, resulting in significantly attenuated *IFIT1* induction upon TLR3 activation (Fig 2J), further emphasizing the role of RBM39 on the TLR3 pathway.

Overall, our results suggest that RBM39 was involved in the direct induction of IFNβ and ISGs upon stimulation with dsRNA, but had a minor influence on NF-κB signaling, suggesting that the IRF3 pathway was mainly affected.

### RBM39 controls the RIG-I/MDA5 pathway

Given that Huh7.5 cells, a subclone of Huh7 (*35*), did not express functional PRRs, ectopic expression of retinoic acid inducible gene I (RIG-I) and melanoma differentiation-associated gene 5 (MDA5) allowed us to identify the function of RBM39 in other key innate immune pathways governed by IRF3 (Fig. 3A) (*28*). Notably, silencing of *RBM39* decreased RIG-I and MDA5-mediated *IFIT1* mRNA expression upon transfection of poly(I:C) (Fig. 3B and C). To validate our finding in non-hepatic cell lines, we tested A549 cells, which are based on adenocarcinoma cells from human alveolar basal epithelium. These cells endogenously express RIG-I and MDA5, but not TLR3, and are regarded as innate immune-competent (*36*). Indeed, knockdown of *RBM39* in A549 cells reduced *IFIT1* induction in response to poly(I:C) similarly as observed for PH5CH and Huh7-Lunet (Fig. 3D). Additionally, Indisulam induced RBM39 degradation also significantly reduced RIG-I/MDA5 responses in A549 cells (Fig. 3E), collectively demonstrating that RBM39’s involvement in innate immunity is neither limited to hepatocytes, nor to TLR3 signaling.

**Fig. 3.**
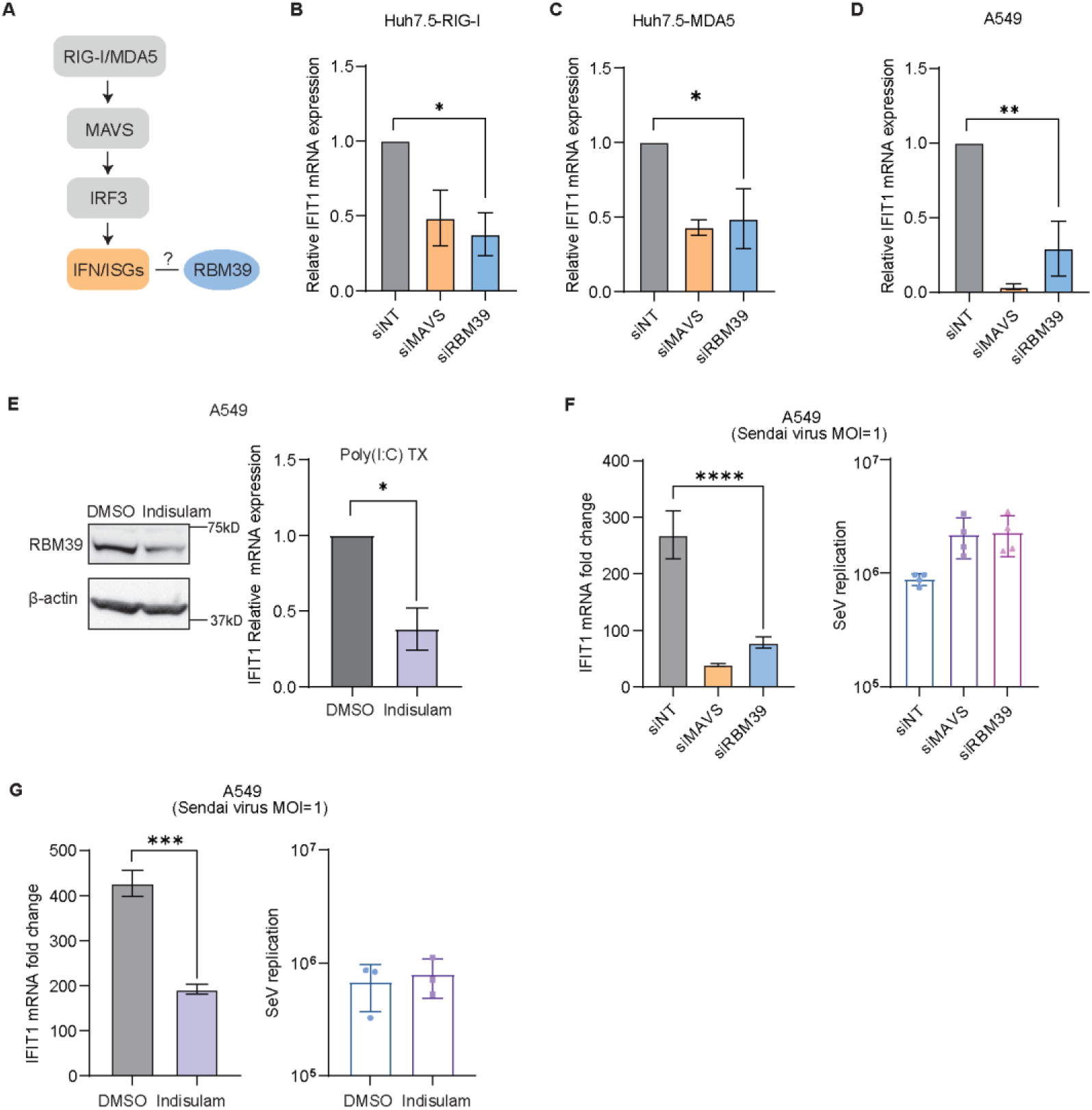
RBM39 participates in the RIG-I/MDA5 response. (A) Simplified schematic of the RIG-I/MDA5-IRF3 pathway. (**B** to **D**) Huh7.5 cells with ectopic RIG-I (B) or MDA5 (**C**) expression, and A549 cells (**D**) were transfected with siRBM39 or siNT/siMAVS as control for 48 h and then transfected with 0.5 µg/ml poly(I:C) for 6 h*. IFIT1* mRNA was measured by RT-qPCR and is shown relative to siNT. (**E**) A549 cells were treated Indisulam or DMSO as control for 48 h, and then transfected with 0.5 µg/ml poly (I:C) for 6 h. RBM39 degradation was determined by immunoblotting. *IFIT1* mRNA was quantified by RT-qPCR. (**F** and **G**) A549 cells were transfected with siRBM39 or siNT/siMAVS as control (**F**) or treated with 1 µM Indisulam or DMSO as control (**G**). 48 hours after treatment, cells were infected with Sendai virus (MOI= 1) for 24 h. *IFIT1* mRNA (left) and Sendai virus P protein mRNA (right) were measured through qPCR. mRNA fold change was normalized on *GAPDH*. Data are obtained from three biological replicates (n = 3), error bars refer to SD. Statistical significance was assessed through Welsch’s unpaired *t*-test.

So far, our study exclusively relied on stimulation of cell intrinsic innate immune responses using the synthetic dsRNA analog poly(I:C). To investigate the function of RBM39 in innate immunity in more physiological settings, we performed virus infection experiments with various viruses known to induce strong innate immune responses mediated by IRF3.

Sendai virus (SeV) is a model virus sensed by RIG-I (*37*). Knockdown of *RBM39* in A549 cells indeed resulted in a strong reduction of *IFIT1* mRNA expression, comparable to the one upon *MAVS* knockdown. We further assessed viral RNA abundance to ensure that RMB39 depletion would not inhibit viral replication, resulting in lower levels of viral stimuli. To this end, we found a slight, comparable increase for both conditions (Fig. 3F). Indisulam was used as an alternative way to delete RBM39 in A549 cells, consistently, it also attenuated SeV-induced *IFIT1* expression, again without affecting viral RNA loads. Similar results were obtained for infection of A549 cells with rift valley fever virus (RVFV) and vesicular stomatitis virus (VSV), HepG2/C3A cells with hepatitis E virus (HEV), and HepG2-NTCP cells with hepatitis D virus (HDV) upon Indisulam treatment, resulting in reduced *IFIT1* mRNA induction, in absence of inhibitory effects on viral RNA replication (Fig. S3A to D).

Altogether, our data demonstrate that RBM39 participated in the regulation of RIG-I and MDA5 pathways and substantially contributed to the cell intrinsic innate immune response to viral infections.

### RBM39 regulates basal IRF3 expression to modulate innate immune responses

In order to elucidate the mechanism involved in RBM39-mediated regulation of IFN and ISG induction, we firstly examined the phosphorylation of IRF3, a key event mediating IFN and ISG induction upon PRR activation by dsRNA. As expected, phosphorylated-IRF3 (p-IRF3) was undetectable in non-stimulated cells and strongly enhanced upon poly(I:C) stimulation. Intriguingly, *RBM39* knockdown not only resulted in a remarkable decrease in p-IRF3, but also in significant reduction of total IRF3 protein levels, suggesting a regulation of the constitutive expression of IRF3 (Fig. 4A).

**Fig. 4.**
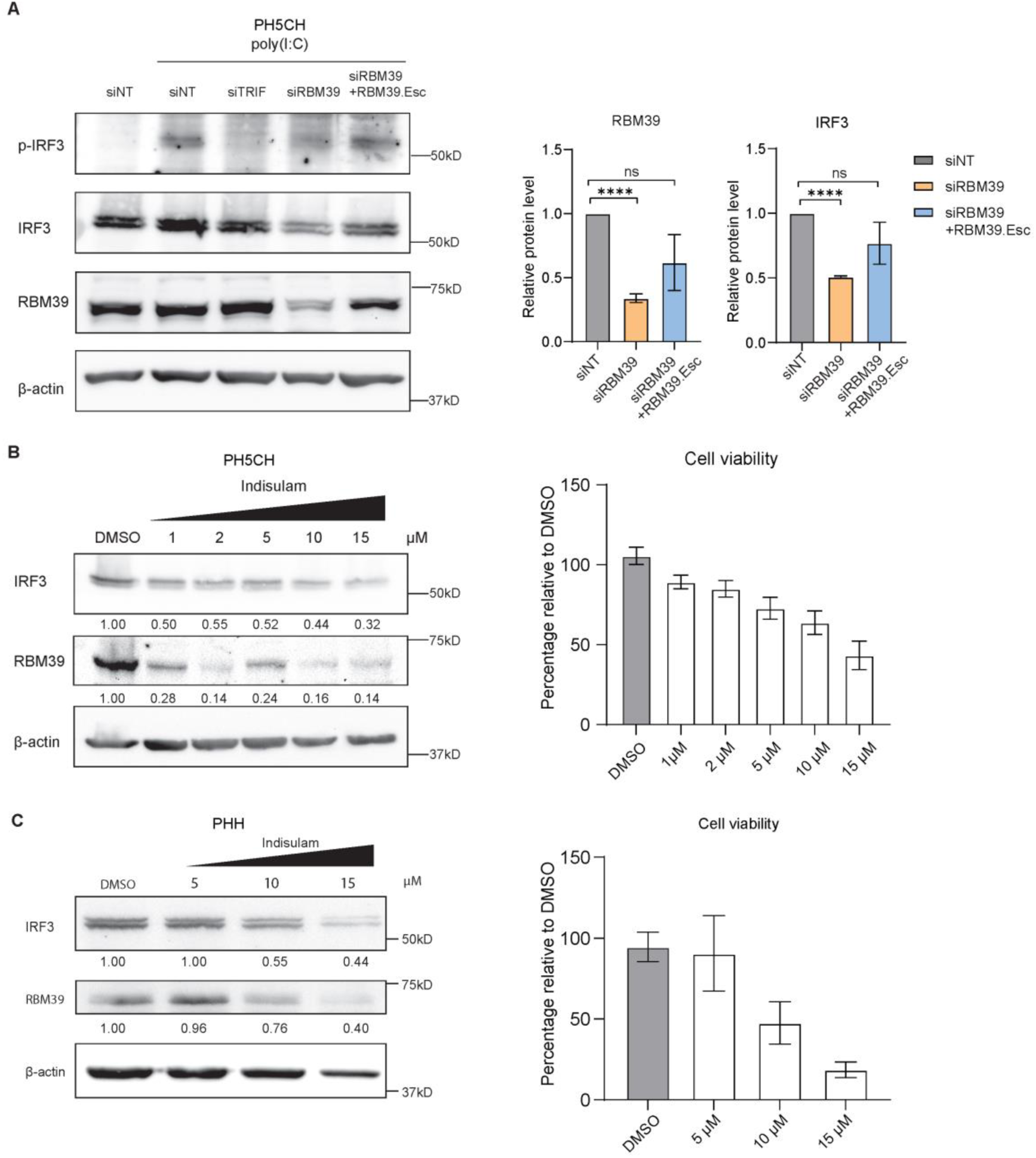
RBM39 controls the basal expression level of IRF3. (**A**) PH5CH cells or PH5CH RBM39.Esc cells were transfected with siRBM39 or siNT/siTRIF as controls for 48 h and then fed with 50 µg/ml poly (I:C) for 6 h. Phosphorylated IRF3 (p-IRF3), IRF3, RBM39 and β-actin protein expression levels were assessed by western blot (left). Quantification of three biological replicates was shown (right). (**B** and **C**) PH5CH cells (**B**) and primary human hepatocytes (PHH) (**C**) were treated with increasing concentrations of Indisulam as indicated. IRF3, RBM39 and β-actin protein expression levels were detected through immunoblotting (left). Cell viability was measured via CellTiter-Glo luminescent cell viability assay (right). Quantification of three independent experiments was expressed as average and is shown under each band. Protein expression was normalized on β-actin, relative protein levels to siNT or DMSO control were shown. Data shown are from three biological replicates (n = 3), error bars indicate SD. Statistical significance was assessed through Welsch’s unpaired *t*-test.

To further validate the impact of RBM39 on IRF3 expression, we treated PH5CH cells with Indisulam. Indeed, RBM39 protein levels were reduced in Indisulam-treated cells in a dose-dependent manner, concomitant with lower IRF3 abundance and some cytopathic effect at higher concentrations (Fig. 4B). In order to verify whether IRF3 depletion upon Indisulam treatment was mediated by RBM39, we made use of an RBM39 mutant (G268V) reported to confer partial resistance to Indisulam (*22*). Overexpression of RBM39 G268V partially rescued RBM39 Indisulam-mediated degradation and thus restored IRF3 expression compared to cells expressing wild-type RBM39 (Fig. S4A). These results confirmed that Indisulam affected IRF3 expression by targeting RBM39.

Similar results were obtained in A549 cells (Fig. S4B) and in primary human hepatocytes (PHH) (Fig. 4C), with the latter indicating that the RBM39-mediated IRF3 depletion also appeared in primary cells, albeit requiring higher concentrations associated with substantial cytotoxicity (Fig. 4C). In contrast, RBM39 was refractory to Indisulam treatment in primary mouse hepatocytes (PMH) (Fig. S4C), probably due to the lack of expression of murine DCAF15 or resistance to Indisulam (*38*), in line with the lack of cytotoxicity (Fig. S4C, right).

Above all, our results demonstrated that RBM39 governs basal expression of IRF3, thereby exerting a crucial role in the activation of innate immune responses.

### Global transcriptome, splicing and proteome analysis highlights the role of RBM39 in innate immunity

We next aimed for a comprehensive proteomic and transcriptomic analysis, to identify other components of innate immunity affected by RBM39, comparing the impact of siRNA-mediated *RBM39* knockdown and Indisulam treatment in PH5CH cells.

For the proteomic analysis we found 165 proteins in *RBM39* knockdown samples and 627 proteins in Indisulam-treated samples with significant changes (fold change < 0.5 or > 2, p-value <0.05) (Fig. 5A and B, Table S2). RBM39 was among the most significantly changed proteins, as expected (Fig. 5A and S5B), whereas GAPDH signals were not significantly changed (Fig. S5A), arguing against a strong impact of cytotoxic effects. IRF3 levels were significantly reduced in both conditions, even though the changes upon RBM39 knockdown were mild (Fig. 5A and B, S5C). The generally lower magnitude of effects upon RBM39 knockdown might be mediated by the difference in the time-frame of RBM39 reduction, acting faster due to direct protein degradation by Indisulam treatment compared to the time required for siRNA-mediated reduction of RBM39 abundance. Importantly, we identified additional factors critically contributing to the cell intrinsic innate immune response, including RIG-I, MDA5, STAT1 and STAT2 (Fig. 5A and B, S5D to G). Similar to IRF3, all these factors showed significant changes in both conditions, with strong reduction in Indisulam-treated samples and mild decline in RBM39 knockdown samples, except for RIG-I that only was significantly decreased in Indisulam-treated samples (Fig. S5E).

**Fig. 5.**
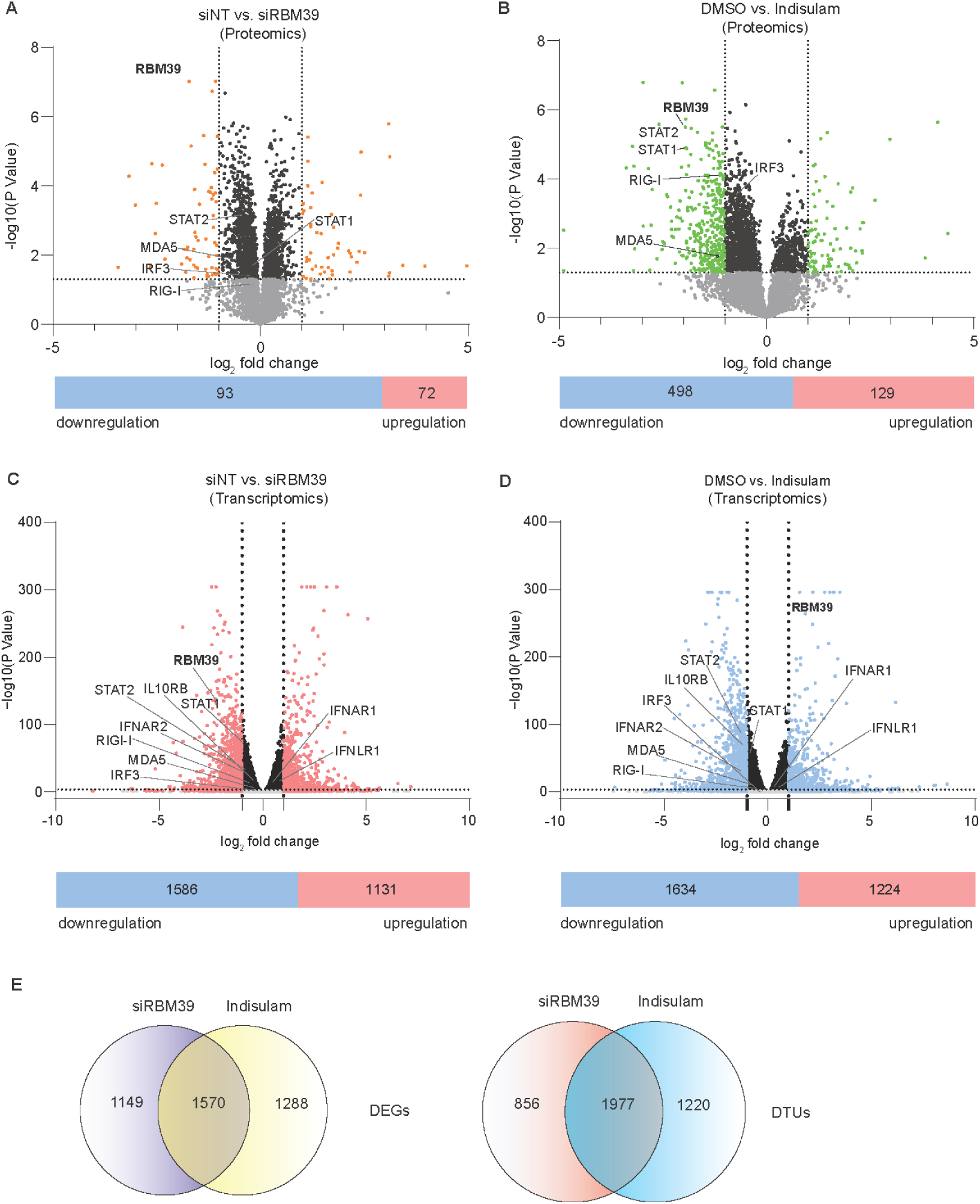
Global proteomic, transcriptomic and splicing analysis upon modulation of RBM39 abundance. PH5CH cells were treated with siRBM39 vs. siNT or Indisulam vs. DMSO after 48 h. The proteome was measured by mass-spectrometry. The transcriptome and splicing were measured through RNA-sequencing. (**A** and **B**) Volcano plot of the proteomic data for RBM39 knockdown (A) or Indisulam-treated (B) samples, the numbers of down- and up-regulated factors were shown under the plot. Factors with a fold change >2 or <0.5 and a p value <0.05 were highlighted with orange (A) or green (B). All factors with a p value <0.05 but lower than 2fold change are shown in black. IRF3, RBM39, RIG-I, MDA5, STAT1 and STAT2 were highlighted. (**C** and **D**) Volcano plot of DEGs in siRBM39-(C) and Indisulam-treated (D) samples, the numbers of down- and up-regulated factors were shown under the plot. DEGs with a fold change >2 or <0.5 and a p value <0.05 were highlighted with red (C) and blue (D). All factors with a p value <0.05 but lower than 2fold change are shown in black. IRF3, RBM39, RIG-I, MDA5, STAT1, STAT2, IFNAR1, IFNAR2, IFNLR1, IL10RB were highlighted. (**E**) RNA-seq data is presented as differential expressed gene (DEG) and different transcript usage (DTU). The overview of DEG and DTU numbers in each condition and overlaps are indicated. RNA-seq data are from three biological replicates (n = 3), proteomic data are from four replicates (n=4). Proteomic data was analyzed via Perseus1.6.15.0. DEG and DTU analysis of individual genes were performed using DESeq2 and DRIMseq, respectively.

Due to the complex, multifactorial mode of action of RBM39 in transcription and splicing, we analyzed the RNAseq dataset in the context of differentially expressed genes (DEG) and differential transcript usages (DTUs), indicative of changes in splicing, comparing knockdown of *RBM39* and Indisulam treatment. Our analysis identified 4,007 DEGs (fold change < 0.5 or > 2, p-value <0.05) and 4,053 DTUs (p-value >0.05) in *RBM39* knockdown and Indisulam-treated samples. 1,570 DEGs and 1,977 DTUs overlapped in both conditions (39% and 49% of total events, respectively) (Fig. 5C-E, Table S3 and Table S4), arguing for the reliability of the analysis. Comparing with proteomic data, DEG analysis revealed more factors with significant fold change and similar numbers as previous transcriptomic studies (Fig. 5C and D, Table S3) (*21, 23, 39, 40*). Interestingly, *RBM39* RNA abundance was significantly reduced upon knockdown (Fig. 5C), as expected, but strikingly increased upon Indisulam treatment due to autoregulation of *RBM39* mRNA levels by protein abundance (Fig. 5D) (*41*). Importantly and in agreement with the proteomic data, we found significant changes in factors involved in the cell intrinsic innate immune response, namely RIG-I, MDA5, STAT1, and STAT2, but in addition, type I and type III IFN receptors.

In summary, proteomic and transcriptomic analysis consistently identified several key factors of the interferon pathway beyond IRF3 that were regulated by RBM39.

### RBM39 governs the transcription and splicing of crucial innate immune factors

Next, we aimed at elucidating whether reduction of mRNA and protein abundance of the key factors of the IFN-response by RBM39 was mediated by alternative splicing or transcriptional regulation.

For IRF3, we found a non-significant 20% reduction *of IRF3* mRNA read counts upon knockdown of RBM39 or Indisulam treatment (Fig. 6A, left), suggesting that transcriptional regulation was not the main mode of action. This was confirmed by analyzing the impact of RBM39 knockdown on *IRF3* mRNA by RT-qPCR (Figure S6A), and on activity of the *IRF3* promoter (*42*) using luciferase reporter plasmids (Fig. S6B, C) both showing a similar magnitude of reduction. In contrast, DTU analysis revealed substantial, consistent changes in splicing of *IRF3* mRNA when RBM39 was suppressed either by siRNA or by Indisulam treatment, with a significant decrease of the functional *IRF3-203* isoform and an increased abundance of the particular isoform *IRF3-228*, which has been reported to have an inhibitory effect on innate immune activation (*43*) (Fig. 6A, right panel). This data was validated upon *RBM39* knockdown via RT-qPCR using primers specifically targeting each of the *IRF3* isoforms that were detected in RNAseq, obtaining very similar results (Fig. S6D). To exclude the contribution of additional factors in the TLR3 pathway potentially affected by RMB39, we expressed IRF3 ectopically in PH5CH IRF3 KO cells. Indeed, overexpression of IRF3 restored the *IFIT1* mRNA induction upon RBM39 knockdown and TLR3 stimulation (Fig. 6B, C), suggesting that IRF3 was the sole determinant of the TLR3 response regulated by RBM39.

**Fig. 6.**
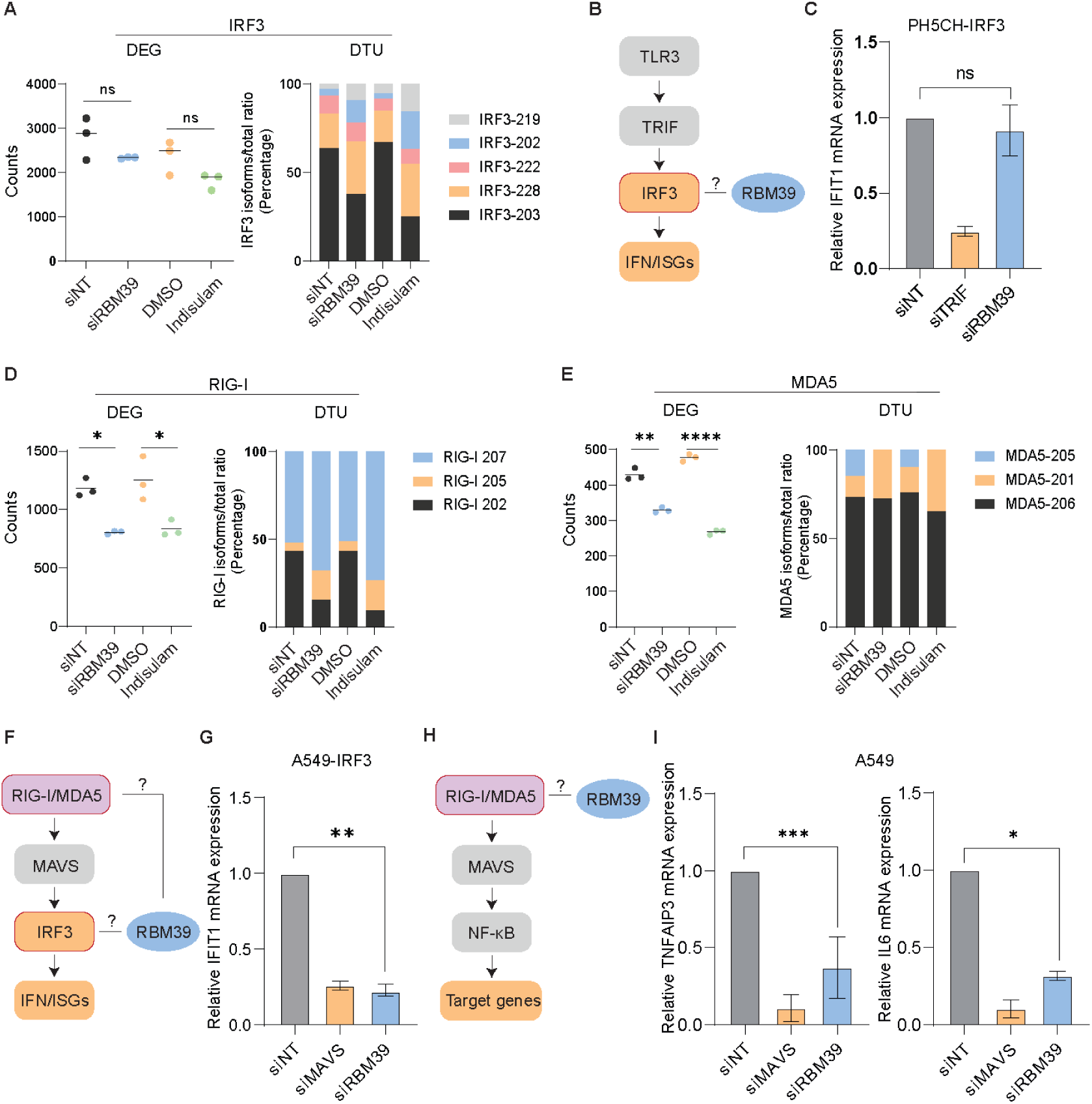
RBM39 controls basal expression of IRF*3* and *RIG-I and MDA5*. (**A**) DEG and DTU analysis of *IRF3* mRNA. (**B**) Simplified schematic of the TLR3 pathway. (**C**) IRF3 was ectopically expressed in PH5CH IRF3 KO cells and cells were transfected with siRBM39 or siNT/siTRIF. 48 h after the knockdown, cells were fed with 50 µg/ml poly (I:C) in the supernatant for 6 h. *IFIT1* mRNA expression levels were measured by RT-qPCR. (**D** and **E**) DEG and DTU analysis of *RIG-I* (**D**) and *MDA5* (**E**) mRNA. (**F**) Simplified schematic of RIG-I/MDA5-IRF3 signaling. (**G**) A549 cells overexpressing IRF3 were transfected with siRBM39 or siNT/siMAVS as controls for 48 h. After the knockdown, cells were transfected with 0.5 µg/ml poly (I:C) in the supernatant for 6 h. *IFIT1* mRNA expression levels were measured by RT-qPCR. (**H**) Simplified schematic of the RIG-I/MDA5-NF-κB pathway. (**I**) A549 cells were silenced with siRBM39 or siNT/siTRIF for 48 h and then transfected with 0.5 µg/ml poly (I:C) for 6 h. *TNFAIP3* and *IL6* mRNA expression levels were measured by RT-qPCR. Data shown are from three biological replicates (n = 3), error bars indicate SD. Statistical significance was assessed through Welsch’s unpaired *t*-test.

In addition to IRF3, the cytosolic dsRNA sensors RIG-I and MDA5 were found to be significantly downregulated in the proteomic and transcriptomic analysis (Fig. 5/S5). DEG/DTU analysis revealed that *RIG-I* expression was modulated both by transcription and splicing regulation, the latter resulting in a lower abundance of the functional isoform 202 (Fig. 6D), whereas *MDA5* expression was mainly decreased at the transcriptional level (Fig. 6E). To verify the consequence of RBM39 regulation on the RIG-I/MDA5 pathways beyond IRF3 (Fig. 6F), we overexpressed IRF3 in A549 cells, thereby excluding any contribution of TLR3. In contrast to the TLR3 pathway in PH5CH cells, IRF3 expression did not suffice to restore *IFIT1* mRNA induction upon RBM39 knockdown and poly(I:C) transfection (Fig. 6G), indicating additional contributions of *RIG-I* and *MDA5*. Consequently, induction of the NF-kB-specific genes *TNFAIP3* and *IL6*, was significantly reduced upon RBM39 knockdown, when the pathway was triggered via RIG-I/MDA5 (Fig. 6H, I).

Taken together, we found that basal expression of IRF3 and RIG-I/MDA5 was regulated by the abundance of RBM39, with significant impact on the induction of cell intrinsic innate immune responses.

### RBM39 regulates the IFN-JAK-STAT pathway

Our proteomic and transcriptomic analysis further identified components of the IFN-JAK-STAT pathway being regulated by RBM39, namely STAT1 and 2 and type I and III IFN receptors, the latter were not detectable by proteomics likely due to low protein abundance. We therefore aimed to understand the mode of regulation and the functional relevance for the IFN response.

The receptor for type I IFNs is a heterodimer consisting of IFNAR1 and IFNAR2. While *IFNAR1* expression was upregulated only on the transcriptional level (Fig. S6E), *IFNAR2* mRNA abundance was reduced and splicing was modulated upon RBM39 knockdown and Indisulam treatment (Fig. 7A). The type III interferon receptor is a heterodimer as well, composed of IFNLR1 and interleukin 10 receptor subunit beta (IL10RB), and also here the expression of both subunits was regulated by RBM39 in opposite directions (Fig. S6F and Fig. 7B), with an increase for *IFNLR1* (Fig. S6F) and a decrease for *IL10RB* (Fig. 7B) RNA amounts, with minor or undetectable changes in the splicing pattern, respectively.

**Fig. 7.**
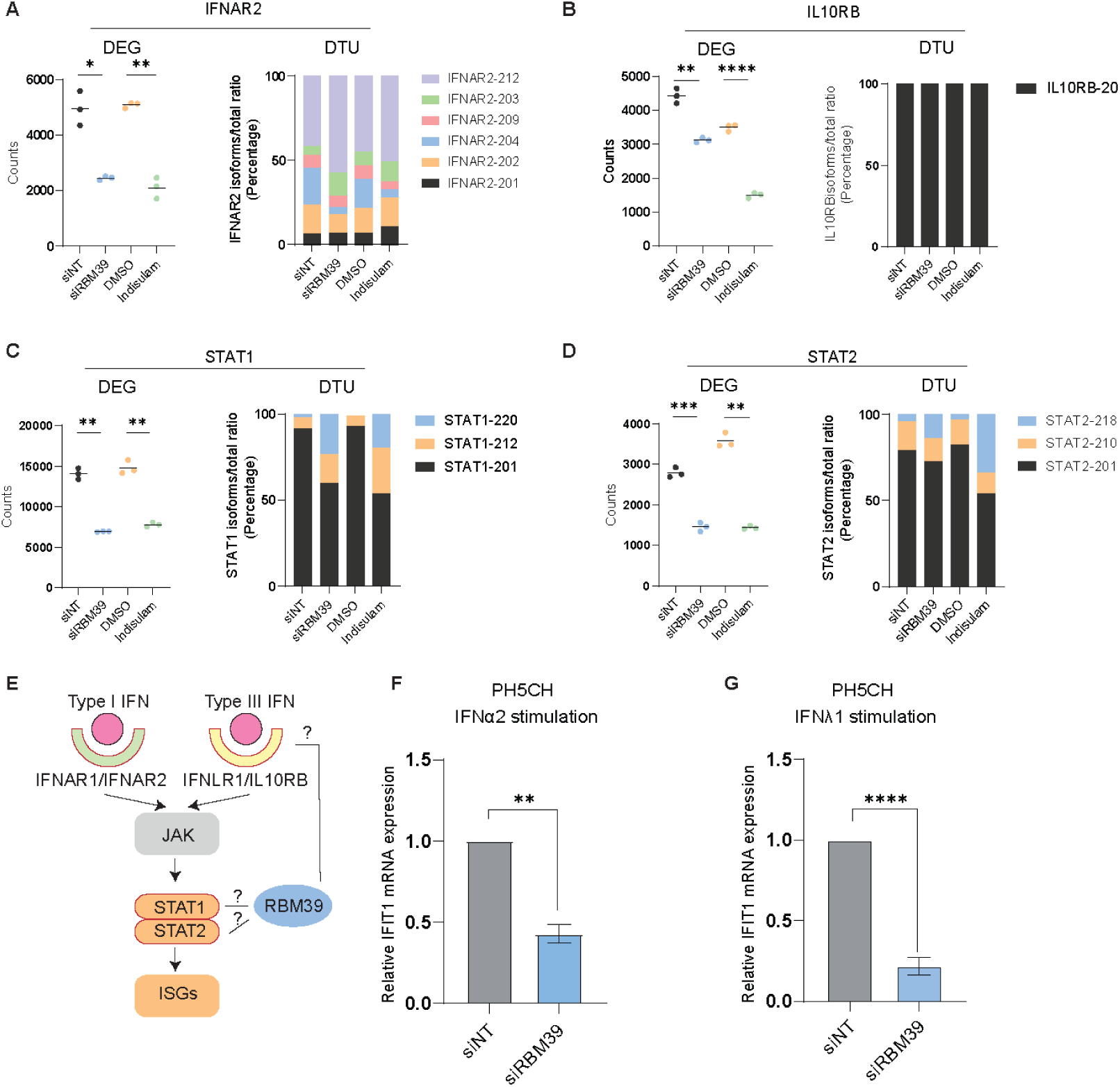
RBM39 impacts on IFN-JAK-STAT signaling. (**A** to **D**) DEG and DTU analysis of *IFNAR2* (**A**), *IL10RB* (**B**), *STAT1* (**C**) and *STAT2* (**D**) mRNA. (**E**) Simplified schematic of the type I and type III IFN pathways. (**F** and **G**) PH5CH cells were transfected with siRBM39 or siNT as control for 48 h and then stimulated with IFNα2 (**F**) or IFNλ1 (**G**) for 24 h. *IFIT1* mRNA was measured by qPCR. Relative expression to siNT control is shown, mRNA fold change was normalized on *GAPDH*. Data are obtained from three biological replicates (n = 3), error bars refer to SD. Statistical significance was assessed through Welsch’s unpaired *t*-test.

STAT1 and STAT2, forming heterodimers upon activation of the type I and type III IFN responses, were in contrast both reduced in protein expression levels upon depletion of RBM39 (Fig. S5F, G), mediated by changes in RNA abundance and associated with modified splicing patterns (Fig. 7C, D).

To investigate the cumulative effect of up- or down-regulation of these factors on the respective IFN response pathways (Fig. 7E), we silenced RBM39 in PH5CH cells and stimulated them with IFNα2 and IFNλ1 to activate type I and type III IFN pathways, respectively. Consequently, the type I and type III IFN responses were both strongly impaired, as indicated by reduced IFIT1 induction (Fig. 7F and G), suggesting that RBM39 also regulates IFN-JAK-STAT signaling.

All in all, our RNA-seq and proteome analysis identified key factors of the IFN response affected by RBM39, such as IRF3, RIG-I, MDA5, STAT1 and STAT2 and type I/III receptors, arguing for a crucial role of RBM39 in the regulation of innate immunity.

## Discussion

Genome-wide CRISPR/Cas9 screens present a powerful tool to study host factors of innate immunity mostly in immune cells, such as macrophages (*44*), dendritic cells (*45*), or in the more commonly used Hela cells (*46*) and HEK 293T cells (*47*). In our study, we applied this approach in two liver cell lines, Huh7 and PH5CH, and specifically targeted the TLR3 pathway by poly(I:C) supernatant feeding. Eventually, we identified RBM39 as a novel factor that affects TLR3 signaling and has a broad impact on the cell intrinsic innate immune response by regulating the basal expression of IRF3 and other important pathway components.

RBM39 is a factor involved in splicing and transcriptional regulation of many genes, particularly contributing to tumorigenesis (*20, 21, 48*), cell cycle control (*12, 19*), and metabolism (*19*), but so far its role in innate immunity has not been shown. Paradoxically, a previous study indicated that porcine reproductive and respiratory syndrome virus (PRRSV) infection upregulated porcine RBM39 expression, resulting in dephosphorylation of c-Jun, and subsequent downregulation of AP-1 signaling (*49*). Additionally, studies have reported that some viruses regulate splicing factors like RBM10 and U5 spliceosomal RNA (snRNA) as an evasion strategy (*50, 51*). While RBM39 has previously been shown to interact with the viral NF-κB homolog v-Rel, inhibiting its activation, the interplay between RBM39 and other human host NF-κB members like p50 and p105 has not been explored. Here, we identified basal expression of *IRF3* as one of the key factors of cell intrinsic innate immune responses being modulated by knockdown of *RBM39* and Indisulam-mediated RBM39 degradation. This resulted in significantly impaired activation upon both dsRNA administration and virus infection. In addition, we identified additional pathway components being modulated by RBM39, including the PRRs RIG-I and MDA5, IL-10RB, IFNAR2, STAT1 and STAT2. These factors all contributed to the reduced activation of the interferon response. Overall, this is supposed to result in an increased level of viral replication. In our infection experiments RBM39 depletion only slightly stimulated viral RNA abundance, most likely due to the limited timeframe, providing not sufficient time to establish more pronounced phenotypes.

The complexity of regulatory functions mediated by RBM39 might explain why other transcriptomic and proteomic analyses missed its role in innate immunity so far (*22, 23, 39, 52–55*). In the transcriptomic and splicing analysis, knockdown of *RBM39* by siRNA and Indisulam treatment showed a wide degree of overlap, arguing for the specificity of Indisulam towards RBM39 in the liver-derived cell lines used in our study and in agreement with previous studies (*53, 56*). Overall, the transcriptomic analysis was in our hands more prone to identify innate immune response factors regulated by RBM39. While we could demonstrate that changes in IRF3, as well as RIG-I/MDA5 expression contributed to the induction of cell intrinsic innate immune response, the data were more complex in the case of IFN treatment. Type I and type III IFNs bind to different receptors, but engage identical signaling pathways (*57*). We observed reduced expression of *IL10RB*, *IFNAR2*, *STAT1* and *STAT2* but increased expression of *IFNAR1* and *IFNLR1*, all essential for signal transduction, with STAT1 and STAT2 being involved in the response pathway to both type I and type III IFNs (*57*). Importantly, we confirmed that the aforementioned changes of these factors in sum resulted in the attenuation of both IFN pathways. In addition, further systemic effects of RBM39 ablation on innate immune responses are possible due to the complex involvement of IL-10RB in several cytokine receptor complexes, responding not only to IFNλ, but also to pro- and anti-inflammatory cytokines including IL-10, IL-22 and IL-28/29 (*58, 59*).

RBM39 depletion was shown to have a strong anti-proliferative impact, which might explain our failure to obtain RBM39 knockout clones in PH5CH cells (not shown). This growth inhibitory effect has put forward RBM39 as a promising target for cancer therapy and contributed to numerous clinical trials implementing aryl sulfonamides like Indisulam, albeit yet with limited success, likely due to the lack of predictive markers (*60*). In contrast, Indisulam showed high activity in preclinical models like mice engrafted with human tumors (*61*) and, more recently, in neuroblastoma models (*23, 61*), associated with limited toxicity. Indisulam treatment of all human hepatic cells tested in our study, including two cancer cell lines (HepG2 and Huh7), one non-neoplastic cell line (PH5CH), and PHH, resulted in a significant degradation of RBM39, albeit at varying doses. Although RBM39 protein expression was similar in all these cell lines, PHH could be less sensitive to Indisulam due to lower *DCAF15* expression levels, critical for its mode of action. Importantly, RBM39 degradation was associated with a certain degree of cytotoxicity in all of our cell culture models. In contrast, Indisulam treatment of PMH had no impact on RBM39 and IRF3 expression and lacked obvious cytotoxicity. The different Indisulam sensitivity of PHH and PMH therefore imposes particular caution in interpreting toxicity data obtained in animal models. This is consistent with data showing only a minor effect of Indisulam treatment on RBM39 expression in different organs of mice, including the liver, with significant changes only in murine kidney and the engrafted human tumors (*22, 40, 61*).

The lack of Indisulam activity in PMH further precluded *in vivo* testing regarding immunosuppressive activities *in vivo*. Currently available aryl sulphonamides might generally not be suitable for clinical use as immunosuppressors, due to their focus on anti-cancer activity. However, substances targeting RBM39 to proteasomal degradation based on alternative E3 ligases beyond DCAF15 might be envisaged, as well as alternative tailored depletion strategies like molecular glues and proteolysis targeting chimeras (PROTACs) (*62*). In light of our data defining RBM39 as a regulator of key components of cell intrinsic innate immunity, it could be regarded as a candidate target to mitigate excessive innate immune responses contributing to viral pathogenesis (*63*) or autoimmune diseases (*64*).

## Materials and Methods

### Cell culture

PH5CH cells were a generous gift from K. Shimotohno. Huh 7.5 cells were a kind gift from Charles Rice. A549 cell were obtained from ATCC, A549 IFNAR/LR KO cells (*65*), Huh7-Lunet cells (*66*), Huh7.5-RIG-I and Huh7.5-MDA5 (*28*), HepG2-NTCP (*67*) have been described before. HepG2/C3A were obtained from ATCC. Primary human hepatocytes (PHH) were purchased from BioIVT and cultured in William E media containing 1% (v/v) Glutamax, 1% (v/v) Non-Essential Amino Acids, 1% (v/v) penicillin/streptomycin (all from Gibco, Germany), 0.2% (v/v) normocin (Invivogen, USA), 2% (v/v) B27 (Gibco, Germany), 1% (v/v) N2 supplement (Gibco, Germany), 100 mM nicotinamide (Sigma-Aldrich, USA), 1.25 mM N-acetylcysteine (Sigma-Aldrich, USA), 10 μM Y27632 (Peprotech, USA), 1 μM A83-01 (Tocris, UK). Cells, unless specifically mentioned, were cultured in Dulbecco’s Modified Eagle Medium (Life Technologies, Darmstadt, Germany) supplemented with 10% fetal bovine serum, nonessential amino acids (Life Technologies), 100 U/mL penicillin, and 100 ng/mL streptomycin (Life Technologies) and cultivated at 37°C and 5% CO2. PH5CH-KDM2A, PH5CH-RBM39.Esc, PH5CH-RBM39 G268V, PH5CH-IRF3, A549-IRF3 cells were kept under selection pressure of 1 mg/mL G418 (Geneticin) (Life Technologies, USA), Huh7-Lunet-TLR3, Huh7.5-RIG-I, HepG2-NTCP cells were maintained under selective pressure of 1 μg/mL puromycin (Sigma-Aldrich, Germany), Huh7.5-MDA5 cells were kept under the antibiotic pressure of 5 μg/mL blasticidin (Sigma-Aldrich, Germany).

### siRNA transfection

The used siRNAs were purchased from Horizon Discovery, UK, unless specifically mentioned. siMAVS was purchased from Sigma Aldrich, Germany. For siRNA transfection in a 6-well plate, cells with 60%-70% confluency were seeded 16 hours prior to the experiment and 1.5 μM siRNA were mixed with 9 μl Lipofectamine RNAiMax (Thermo Fisher Scientific, USA) in OptiMEM according to manufacturer’s instructions. After 20 min incubation at RT, the mixture was added into the medium. Cells were incubated for 48 h and used for further experiments.

### Drug treatments

Poly(I:C) stimulation: Poly (I:C) (HMW) was used (Invivogen, USA) following the manufacturer’s instructions. To specifically trigger TLR3, 50 μg/ml poly (I:C) were added directly into the medium. For TLR3, RIG-I and MDA5 stimulation, transfection was performed with 0.5 μg/ml poly (I:C) mixed with Lipofectamine2000 (Thermo Fisher Scientific, USA) in OptiMEM (Gibco, USA) according to the manufacturer’s instructions. After 20 min incubation at room temperature (RT), the mixture was added into the medium. Cells were stimulated for 6 h or 8 h, then harvested for RNA or protein extraction.

Indisulam treatment: Indisulam was utilized (Sigma Aldrich, Germany) solved in DMSO at a final stock concentration of 10 mM. Indisulam, or same amount of DMSO as control, were added to the cell medium at the indicated concentrations; cells were incubated for the indicated time and treated further with either poly (I:C) or virus infection. At the indicated time points, cells were harvested for RNA extraction or protein analysis. IFNs stimulation: IFNα2 and IFNλ1 were purchased from PBL, USA and Peprotech, USA, respectively. 50 IU/ml IFN2α and 10ng/ml IFNλ1 were added to the medium. Cells were stimulated for 24 h and collected for RNA extraction.

### Statistical analysis

Independent biological replicates are denoted with n-numbers. To test for significance, two-tailed unpaired Welch’s test, were performed using GraphPad Prism 8 software (GraphPad Software, La Jolla, CA, USA). ∗p <0.05; ∗∗p <0.01; ∗∗∗p <0.001; ∗∗∗∗p <0.0001.

## Acknowledgments

The authors thank Rahel Klein, Ulrike Herian for excellent technical assistance. We thank Charles M. Rice for Huh7.5 cells and Kunitada Shimotohno for PH5CH cells. We also need to thank the Genomics and Proteomics Core facility of the German Cancer Research Centre for the RNA-seq sample measurement, as well as Dr Julia Mergner, Proteomics Core Facility of the TranslaTUM. Additionally, we thank Zhenfeng Zhang, Julia Welsch, Jun-gen Hu, Zhaozhi Sun for providing reagents, and Cong Si Tran and Minh-Thu Pham for helpful discussions.

## Funding

This work was supported by the Deutsche Forschungsgemeinschaft (DFG, German Research Foundation) – project number 272983813 – TRR 179, to VL, MB, AP, DW, RB, MS, SU and VLDT. VL received additional DFG funding, project number 519777725. TL was supported by the China Scholarship Council. MB and VGM were supported by DFG 236528794. NAR is a holder of a Georg Forster Research Fellowship for Experienced Researchers from the Alexander von Humboldt Foundation.

## Data and materials availability

Details related to materials and methods are provided in the Supplementary material. Transcriptomic data were deposited in the Sequence Read Archive (SRA) under accession PRJNA1028350. Further data supporting the findings of this study is available upon request. Please contact the corresponding author.

## Supplementary Materials

Supplementary materials and methods

Fig. S1: Efficiency of the tBID death reporter and the rescue of KDM2A

Fig. S2: Rescue of TLR3 response by ectopic RBM39 expression in Huh7 Lunet-TLR3 cells

Fig. S3: Indisulam treatment in different virus infection models

Fig. S4: Impact of Indisulam-mediated RBM39 degradation on IRF3 in different cell lines

Fig. S5: Analysis of individual protein expression levels from proteomics data

Fig. S6: The transcription and splicing of IRF3, IFNAR1 and IFNLR1 mRNA

Table S1 Candidate list in CRISPR/Cas9 screen

Table S2 Proteomic analysis of mass spectrometry

Table S3 DEG analysis of RNA seq

Table S4 DTU analysis of RNA seq

Table S5 Primers and antibodies

## Supplementary materials and methods

### Primary mouse hepatocytes isolation

C57BL6/N wild type mice (9-10 weeks, male) were housed in the specific-pathogen free (SPF) barrier at the Interfakultäre Biomedizinische Forschungseinrichtung (IBF) animal facility at Heidelberg University (Germany). Mice are maintained on a standard diet with ad libitum access to food and water under a constant light-dark cycle. Animal experiment was approved by the Regierungspräsidium Karlsruhe (T47/22). Mice were sacrificed, the abdomen was cleaned with ethanol and then opened expose inferior cava vein and portal vein. The inferior cava vein was Cannulated and washed with pre-warmed Liver Perfusion Medium (ThermoFisher, USA) for about 3-4 minutes until the liver became blood free. Afterwards, the liver was perfused with Liver Digest Medium (ThermoFisher, USA) until it was digested. The liver without gallbladder was transferred to a Petri dish. Cold Hepatocyte Wash Medium (ThermoFisher, USA) was added and the organ liberating cells were de-capsuled. Subsequently, the cells were sequentially passed through 100 µm and 70 µm cell strainers into a 50 mL falcon tube and centrifuged at 50g, 4°C for 2 minutes, to separate liver non-parenchymal cells (NPC) that remain in the supernatant. Next, hepatocytes (HC) pellet was washed with Hepatocyte Wash Medium (ThermoFisher, USA) once and re-suspended in William’s E Medium, Glutamax Supplement (ThermoFisher, USA) with 4% FBS, 1% pen-strep and seeded 5 × 10^4^ cells/cm^2^ in a collagen-coated plate (Collagen I Rat Protein, Tail, ThermoFisher, USA).

### CRISPR/Cas9 knockout screening

For genome-wide CRISPR/Cas9 screen, PH5CH and Huh7-Lunet-TLR3 cells were transduced with lentiviral vectors containing the tBID death reporter and Cas9 (lentiCRISPRv2). Neomycin and puromycin were added for selection. After that, PH5CH and Huh7-Lunet-TLR3 cells stably expressing the tBID death reporter and Cas9 were transduced with lentiviral vectors containing the GeCKOv2.0 human genome-wide sgRNA library (Addgene, USA) at MOI=0.3. The GeCKOv2.0 human genome-wide sgRNA library includes a total of 122,411 unique sgRNAs targeting 19,050 genes (with 6 sgRNAs per gene) and 1,864 miRNAs (with 4 sgRNAs per miRNA) and a total number of 1,000 different non-targeting sgRNAs as controls.

After 24 hours, puromycin was added for selection. On day 7 post-transduction, 50ug/ml poly (I:C) or PBS as control were added to the medium to exclusively activate the TLR3 pathway. The experiments were performed in 2 independent biological replicates with 2 technical replicates each. Surviving cells were collected at 72 h post-stimulation, genomic DNA was extracted using NucleoSpin Blood L kit (Macherey-Nagel, Germany) according to manufactureŕs instructions. Amplicons were amplified, barcoded by PCR, and purified again via agarose gel separation using NucleoSpin Gel and PCR Clean-up kit (Macherey-Nagel, Germany), before being pooled and submitted to GATC Biotech (Germany), for Illumina NGS (Illumina HiSeq, 50 bp single reads, 240 million reads total, 20% PhiX DNA).

NGS data first was de-multiplexed to allocate the respective reads in the pooled population to each sample. Reads were counted for each gRNA in every sample, and then normalized to the total number of reads in this sample as well as the complexity sample taken one day after transduction of the lentiviral library, yielding normalized read counts.

Mean values of normalized read counts of the 4 technical replicates were generated for each of the 4 experiments. Based on mean values, fold changes were calculated between poly(I:C)- and PBS-treated samples. gRNAs with a higher fold change than a threshold were considered as primary hits. The threshold was adjusted for each cell line to yield the about 2000 top-ranked gRNAs for each experiment. Thresholds were 2.5x change in Huh-7-Lunet-TLR3 cells and 7x change in tPH5CH cells. Preliminary hits were furthermore filtered for occurrences of at least 5 upregulated gRNAs per gene in all datasets combined, and the occurrence of at least one gRNA per gene in at least 3 independent datasets. This yielded 50 candidate genes and 5 candidate miRNAs.

### CRISPR/Cas9 knockout clones

To generate PH5CH IRF3 single knock-out (KO) clones, PH5CH cells were transduced with the lentiCRISPRv2-IRF3 and lentiviral vectors encoding for Cas9 (see “Plasmids”). After puromycin selection, cells were seeded into 96-well plates for screening of stable single clones.

### Real time quantitative PCR

Total RNA was isolated from cultured cells using the NucleoSpin RNA Plus kit (Macherey-Nagel, Germany). RT-qPCR was performed as described before (*28*). For gene expression analysis, cDNA as generated from RNA samples using High-Capacity cDNA Reverse Transcription Kit (Thermo Fisher Scientific, USA), and was used for qPCR analysis with the 2x iTaq Universal SYBR Green Supermix (Bio-Rad, Germany). Reactions were performed on a CFX96 Touch Real-Time PCR Detection System (Bio-Rad, Germany) as follows: 95°C for 3 minutes, 95°C for 10 and 60°C for 30 seconds. mRNA fold change was calculated normalizing on Glyceraldehyde-3-phosphate dehydrogenase (GAPDH) mRNA expression relative to untreated / uninfected / mock transfected cells. For IRF3 isoforms mRNA measurement, cDNA was prepared by reverse transcription kit (Thermo Fisher Scientific, USA) using oligo(dT)18 primers (Thermo Scientific, Germany). mRNA expression fold change was normalized on GAPDH and Hydroxymethylbilane synthase (HMBS) expression levels relative to siNT-transfected cells. Primers IRF3-228+203 were designed to target both IRF3-228 and IRF3-203 stretches, other IRF3 primers were designed to target specific exons or introns of individual isoforms. IRF3-203 mRNA expression was calculated by subtracting the expression of IRF3-228 from that of IRF3-228 + IRF3-203. All qPCR primers are listed in Table S5.

### Western blot

Cells were lysed in protein lysis buffer (50mM TRIS HCl pH7.4, 150mM NaCl, 1% Triton-X 100, protease inhibitor) on ice for 45 min. Cell lysates were then centrifuged at 14,000 g for 15 min at 4 °C to exclude cell debris. Supernatants were collected and mixed with 6x Lämmli buffer (0.375M Tris-HCl, 9% SDS, 50% Glycerol, 0.03% Bromophenol Blue, 9% β-mercaptoethanol), boiled at 95°C for 5 min and cooled down on ice for 2 min. Samples were separated by 10% SDS-PAGE and then blotted onto polyvinylidene difluoride (PVDF) membranes (Merk Millipore, Germany). Membranes were then blocked with 5% milk in PBST (0.05% Tween) or 5% BSA in TBST (0.01% Tween) and incubated with the indicated antibodies. Membranes were imaged via the ECL Plus Western Blotting Detection System (Pierce, GE Healthcare, Little Chalfont, UK) according to the instructions of the manufacturer. Signal was recorded using the Advance ECL Chemocam Imager (Intas Science Imaging, Germany). Quantification of western blot was performed by Fiji.

### Lentivirus preparation and transduction

5 x 10^6^ HEK293T cells were seeded in 10 cm dishes 16 h before transfection. The next day, the target plasmid (5.14 µg) together with pCMV-Gag-Pol (5.14 µg) and pMD2-VSVG (1.71 µg) were mixed in 400µl OptiMEM (Gibco, USA). 364 µl of OptiMEM were separately mixed with 36µl polyethylenimine (PEI). The two mixtures were merged and incubated for 20 min at RT, and then added to the cells as previously described (*68*). After 2 days, lentivirus was collected and filtered at 0.45 µm filter (Sigma Aldrich, Germany) to exclude cell debris, and store at −80 °C or used for transduction. PH5CH-IRF3, A549-IRF3 stable cells were produced by transduction of lentiviral particles encoding for IRF3 under control of the Rosa26 promoter (IRF3 sgRNA resistant mutant) as previously described (*28*) in the abovementioned PH5CH IRF3 KO cells. PH5CH-KDM2A, PH5CH-RBM39.Esc, PH5CH-RBM39 G268V were generated by lentiviral transduction of particles containing corresponding plasmids.

### Cell Viability Assay

Cell viability upon transduction of tBID death reporter was assessed in 96-well plates through WST-1 assay (Roche, Switzerland) according to the manufacturers’ instructions. Cell viability of Indisulam-treated cells was accessed by CellTiter-Glo® 2.0 Assay (Promega, Germany) according to the manufacturer’s instructions.

### ELISA

Supernatants were collected from stimulated cells, IFNβ protein levels were measured by LumiKineTMXpress hIFN-β 2.0 (InvivoGen, Germany) according to the manufacturer’s instructions.

### Luciferase reporter assay

pGL3-982 or pGL3-149 Firefly reporter, together with pRL-CMV Gaussia (Promega, USA) reporter as reference were transfected into PH5CH RBM39 knockdown cells using Lipofectamine2000 (Thermo Fisher Scientific, USA) according to the manufacturer’s instructions. After 24 h, cells were lysed by addition of Luciferase lysis buffer (1% Triton X-100, 25mM Glycyl Glycin, 15mM MgSO4, 4mM EGTA, 10% Glycerol, pH7.8, 10mM DTT) and luciferase activity (Relative Ligh Units, RLU) was measured by a Mitras LB940 luminometer (Berthold Technologies, Germany). RVFV-RLuc assay was performed as described before (*69*). At 24 hours after infection, infected cells were washed once with PBS and immediately lysed with luciferase lysis buffer. Cell lysates were stored frozen at −80°C to ensure proper lysis and the Renilla luciferase signal was measured by injecting 300 µl of luciferase assay buffer (15 mM K_3_PO_4_ [pH7.8], 25 mM Glycyl-Glycin [pH 7.8], 15 mM MgSO_4_, 4 mM EGTA) supplemented with 1.43 coelenterazine (102173, PJK, Germany) followed by read-out of the signal with a 480 nm high-sense filter (480m20BREThs, Berthold) using the Mitras2 multimode plate reader (LB942, Berthold).

### RNA sequencing (RNA seq)

RNA-sequencing and RNA quality control was performed by the Genomics and Proteomics Core facility of the German Cancer Research Centre using the Illumina Hiseq NovaSeq 6000 with RNA seq reads in 2 x 100 bp format. For sample preparation, 1.5 x 10^5^ cells were seeded in a 6-well plate. Cells were either transfected with siRBM39 or siNT as control, or treated with 1mM Indisulam or the same amount of DMSO as control. 48 hours after transfection, cells were lysed and total RNA was extracted via NucleoSpin RNA Plus kit (Macherey-Nagel, Germany) following the manufacturer’s instructions. RNA was quantified using OD measurement at 260 nm.

Human reference transcriptome (Human Gencode v40) mapping and quantification of the transcripts were performed using the salmon package with GC bias correction (*70*). Differential expression analysis was performed using DESeq2 (*71*), and DTU analysis was performed with DRIMseq (*72*).

### Virus infection

For Sendai virus (SeV) infection, 6 x 10^4^ A549 cells were transfected with siRNA RBM39 or treated with 1 µM Indisulam/DMSO. After 48 h, the cells were infected with SeV (MOI=1) in PBS with 0.3% BSA at room temperature (RT). One hour after infection, cells were washed with phosphate-buffered saline (PBS) and complete medium was added. Cells were collected after 24 h for further analysis.

For vesicular stomatitis virus (VSV) infection, 6 x 10^4^ A549 cells were treated with 1 µM Indisulam or DMSO control for 48h. Afterwards, cells were infected with recombinant VSV-G (rVSV-G) (MOI=0.1) for 24 h. Cells were collected for further analysis.

For Rift Valley Fever Virus (RVFV) infection, 2 x 10^4^ A549 cells were seeded into 24-well plates. The following day, cells were treated with 0.1 µM Indisulam or DMSO for 48 hours, and then inoculated with RVFV-RLuc at a MOI of 0.01 in DMEM with 2% FCS. After 24 hours of infection, cells washed once with PBS, lysed in 100 µl of luciferase lysis buffer, and frozen at −80 °C for further analysis.

For hepatitis E virus (HEV) (Genotype 3 Kernow C1 P6 accession number: JQ679013, a kind gift from Suzanne Emerson, NIH) (*73*) and hepatitis D virus (HDV) infection, 2.5 x 10^4^ HepG2/C3A cells and HepG2-NTCP cells, respectively, were seeded in collagen (Corning, USA)-coated plates and treated with 0.1 µM Indisulam or same amount of DMSO for 48h. Subsequently, cells were infected with HEV (MOI=4) in MEM with 10% FBS or HDV (MOI=4) in DMEM with 1.5% DMSO, 4% PEG. 24 hours after infection, cells were washed with PBS. Cells were incubated for 5 days and collected for further analysis.

### Mass spectrometry (MS)

PH5CH cells were treated with siRBM39 or Indisulam for 48 h, siNT and dimethyl sulfoxide (DMSO) were used as control. Afterwards, cells were harvested and lysed in 300 µl sodium dodecyl sulfate (SDS) lysis buffer (4% SDS, 10 mM dithiothreitol (DTT), 55 mM iodoacetamide (IAA), 50 mM Tris/HCl pH 7.5), and heat-inactivated (95°C for 10 min). The samples were sonicated (4°C, 15x 30 sec on and 30 sec off setting) and the protein lysate was precipitated using pre-chilled acetone (−20°C) at a final concentration of 80% acetone (v/v) overnight. Precipitates were pelleted and washed two times with pre-chilled 80% acetone (−20°C). Pellets were air dried at room temperature and resolubilized by adding 40 µl thiourea buffer (6 M urea, 2 M thiourea (U/T) in 10 mM HEPES, pH 8.0). Protein concentration was determined by BCA assay (Pierce) according to the manufacturer’s instructions and adjusted to 50µg with thiourea buffer. Lysates were digested overnight (25°C, 16h, 1200 rpm) using LysC (1:50 w/w, enzyme/protein, Wako) and trypsin (1:50 w/w, enzyme/protein, Promega) per replicate. The samples were prepared for C18-stage tipping by adding 1/5 of loading buffer (10% acetonitrile (ACN), 3% trifluoroacetic acid (TFA)) to inactivate proteases (TFA) and enhance solubility (ACN) of samples. Peptides were desalted with 3-layer C18 stage tips (3M). Columns were equilibrated with 200 µl methanol (RT, centrifuge 300xg, 4min), washed twice with 200 µl washing buffer (0.5% acetic acid (AA)), loaded with the sample, washed twice with 200 µl washing buffer (0.5% AA) and eluted (80% acetonitrile, 0.5% AA). All steps were executed with 500 g at RT. The eluate was evaporated (45°C, vacuum) and resuspended in MS buffer (2% acetonitrile, 0.3% TFA). Peptide concentrations were measured at 280 nm (Nanodrop 2000, Thermo Scientific), and subsequently equalized to 0.25 µg/µl. Peptides were loaded onto a 2 cm trapping column (75 µm diameter; ReproSil-Pur C18-AQ 5 µm resin; Dr. Maisch) at 5 ul/min 0.1% formic acid (FA) in water using the UltiMate 3000 RSLCnano System (Thermo Fisher). Peptides were eluted onto a 42 cm analytical column (75 µm diameter; ReproSil-Pur C18-AQ 1.9 µm resin; Dr. Maisch) over a 120 min gradient at a flowrate of 300 nl/min and the column oven operating at 50 °C, using solvent B (0.1% FA, 5% DMSO in acetonitrile) starting from 4% to 32% over 110 min, then to 80% over 2 min, and kept at 80% for other 2 min, to 2% over 2min, and kept at 2% for 3 min. nanoLC solvent A was 0.1% FA, 5% DMSO in HPLC grade water.

Spectra were acquired with an Orbitrap Eclipse Tribrid mass spectrometer (Thermo Fisher Scientific) running Tune 3.5 and Xcalibur 4.5, and equipped with a nano-electrospray source (Thermo Fisher Scientific). Spray voltage was set to 2.1 kV, RF lens at 40 %, and heated capillary at 275 °C. The Eclipse mass spectrometer was operated in data-independent acquisition (DIA) and positive ionization mode. MS1 full scans (360–1300 m/z) were recorded at a resolution of 120,000 (Orbitrap) using a normalized automatic gain control (AGC) target value of 100% and maximum injection time of 50 ms. Peptides were fragmented using higher energy collision induced dissociation (HCD) collision energy set to 30%. MS2 scans (200–1800 m/z) were acquired over a total of 40 DIA segments with variable isolation windows, and with a window overlap of 1 m/z. The scan resolution in the Orbitrap was set to 30,000 with a normalized AGC target value of 1000% and a maximum injection time of 54 ms.

Raw MS data files of the experiments conducted in data-independent acquisition (DIA) mode were processed with DIA-NN version 1.8.1 (PMID 31768060). An in-silico predicted spectral library composed of the human proteome (UniprotKB reference proteome, UP000005640, download Okt 2021) and common contaminants (MaxQuant contaminants.fasta) with trypsin as digestion enzyme and one missed cleavage specified was generated in DIA-NN. Subsequently the acquired raw files were processed in library-free mode using DIA-NN default settings and the match between runs function enabled. The R package iq was used to generate MaxLFQ protein abundance output with a Lib.Q.Value and Lib.PG.Q.Value filter setting of 0.01 (PMID 31909781). The ProteinGroups output file was processed with Perseus (version 1.6.15.0). The intensities were expressed as Log2(x), imputed (replacing missing values from normal distribution, with.0.3; down shift 1.8), and annotation included. Statistical significance evaluated by “two-sample tests” (Student’s T-test with permutation-based FDR 0.05 and 250 number of randomizations).

---

**Fig. S1.**
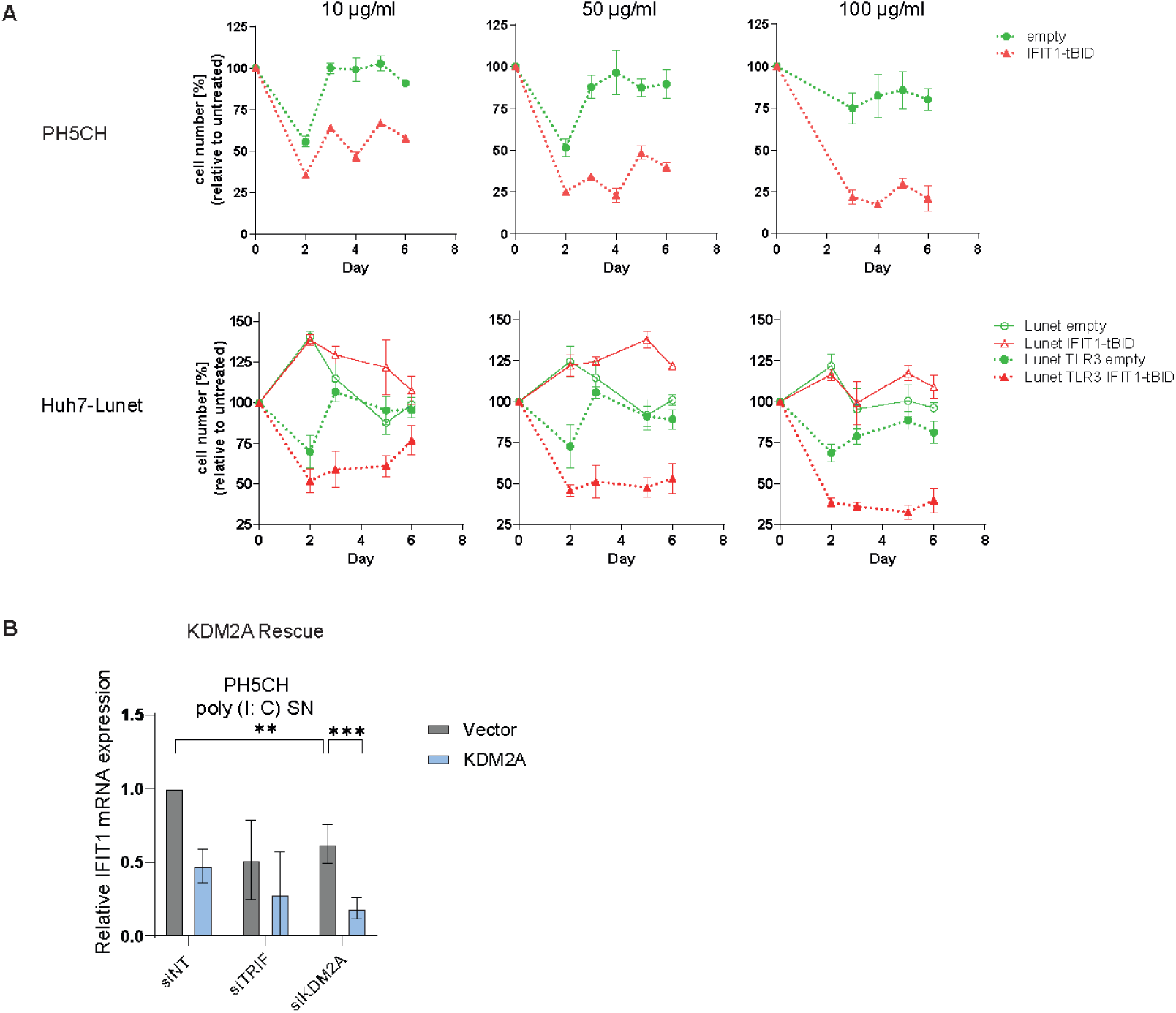
Efficiency of the tBID death reporter and the rescue of KDM2A. (**A**) PH5CH cells (upper panel) or Huh7-Lunet cells, with or without ectopic TLR3 (lower panel), stably expressing IFIT1-tBID or empty-tBID death reporter as control were supernatant-fed with the indicated concentrations of poly(I:C), or a same volume of PBS, for the indicated time. Cell viability was determined at the indicated time points using the WST-1 assay. Data are from four independent biological replicates (n=4). (**B**) PH5CH expressing empty vector or *KDM2A* were transfected with the indicated siRNAs for 48 h. Cells were subsequently stimulated through poly(I:C) supernatant feeding. *IFIT1* mRNA was measured by qPCR. Data are from two biological replicates (n=2), error bars represent SD.

**Fig. S2.**
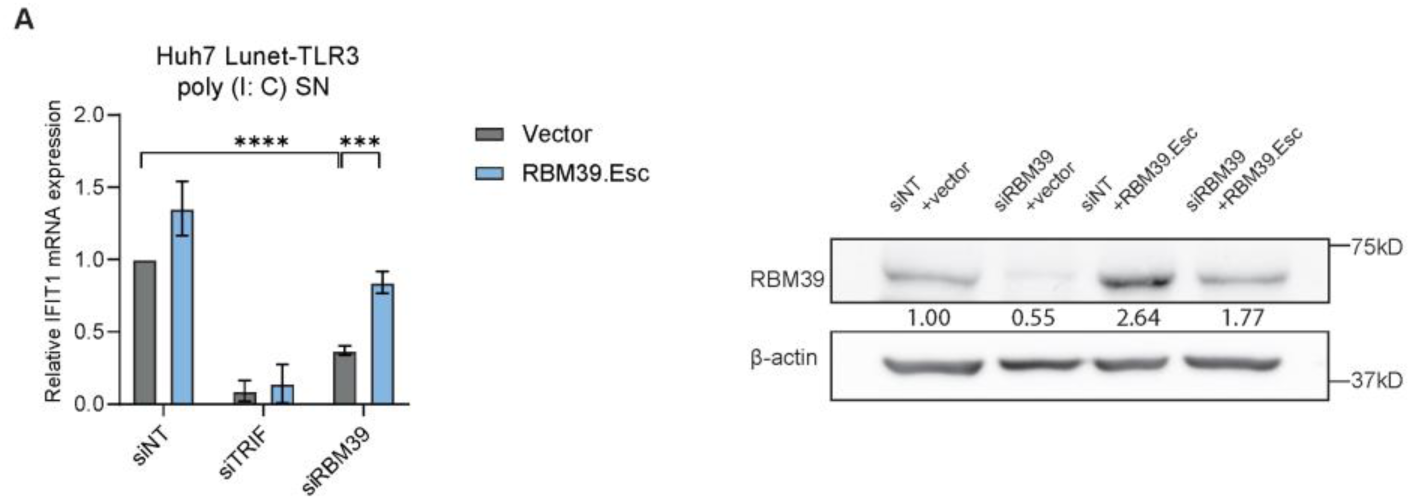
Rescue of TLR3 response by ectopic RBM39 expression in Huh7 Lunet-TLR3 cells. (**A**) Huh7 Lunet-TLR3 cells expressing RBM39.Esc or empty vector were transfected with siRBM39 or siNT/siTRIF as controls for 48 h and then fed with 50 µg/ml poly (I: C) in the supernatant for 6 h. *IFIT1* mRNA was measured by RT-qPCR (left), RBM39 expression was measured by western blot (right). Data are from three biological replicates (n = 3), error bars refer to mean ± SD. Statistical significance was assessed through Welsch’s unpaired *t*-test.

**Fig. S3.**
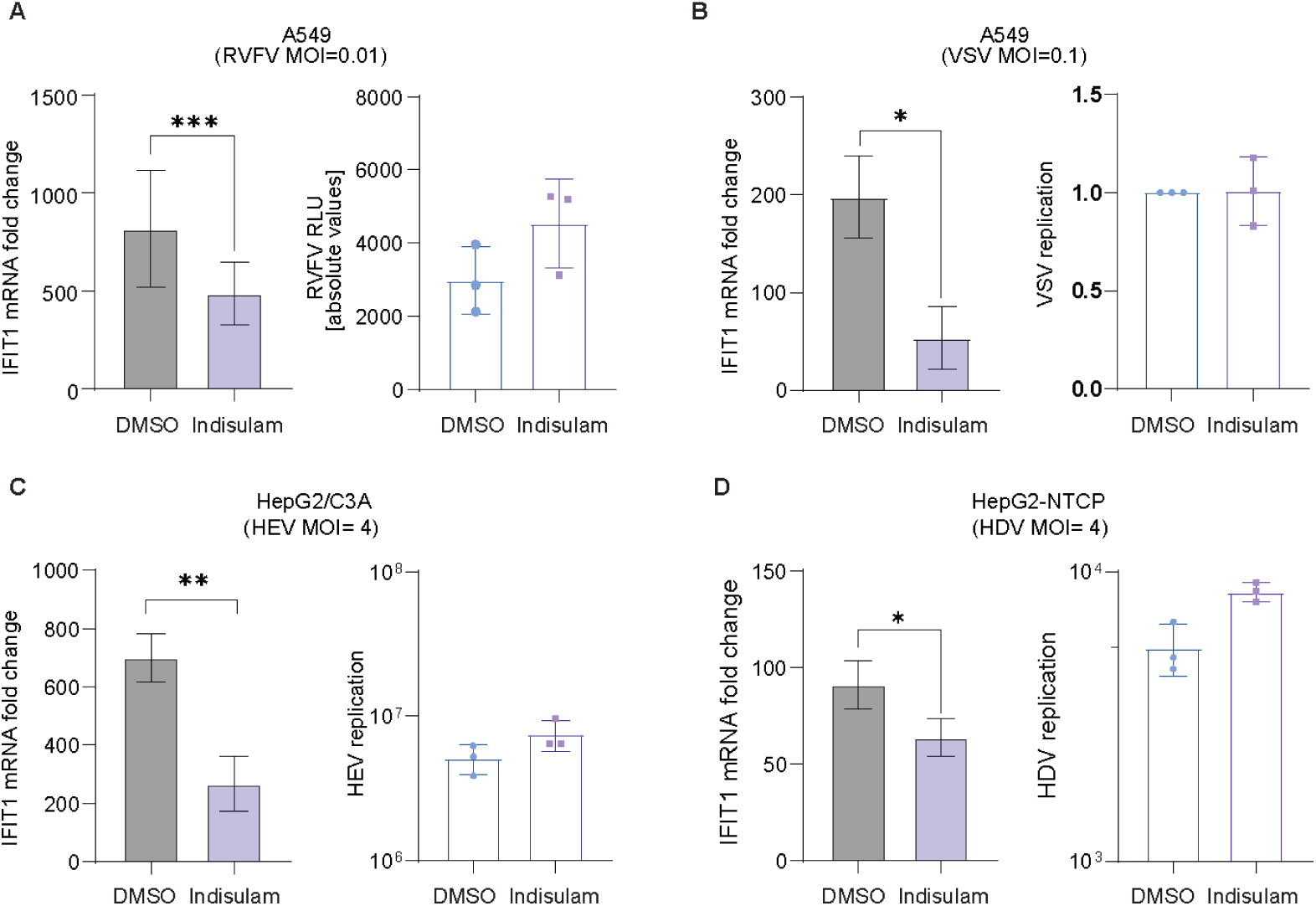
Indisulam treatment in different virus infection models. (**A** to **D**) A549 cells (**A** and **B**), HepG2/C3A (**C**) cells and HepG3-NTCP cells (**D**) were pre-treated with Indisulam or same amounts of DMSO as control for 48 h, and subsequently infected with Luc-RVFV (MOI= 0.01) for 24 h (**A**), rVSV-G (MOI=0.1) for 24 h (**B**), HEV (MOI= 4) (**C**) or HDV (MOI=4) (**D**) for 5 days, respectively. *IFIT1* mRNA (left) and viral mRNA (right) were measured by qPCR. mRNA fold change was normalized on *GAPDH*. Data are derived from three biological replicates (n = 3), error bars refer to SD. Statistical significance was assessed through Welsch’s unpaired *t*-test.

**Fig. S4.**
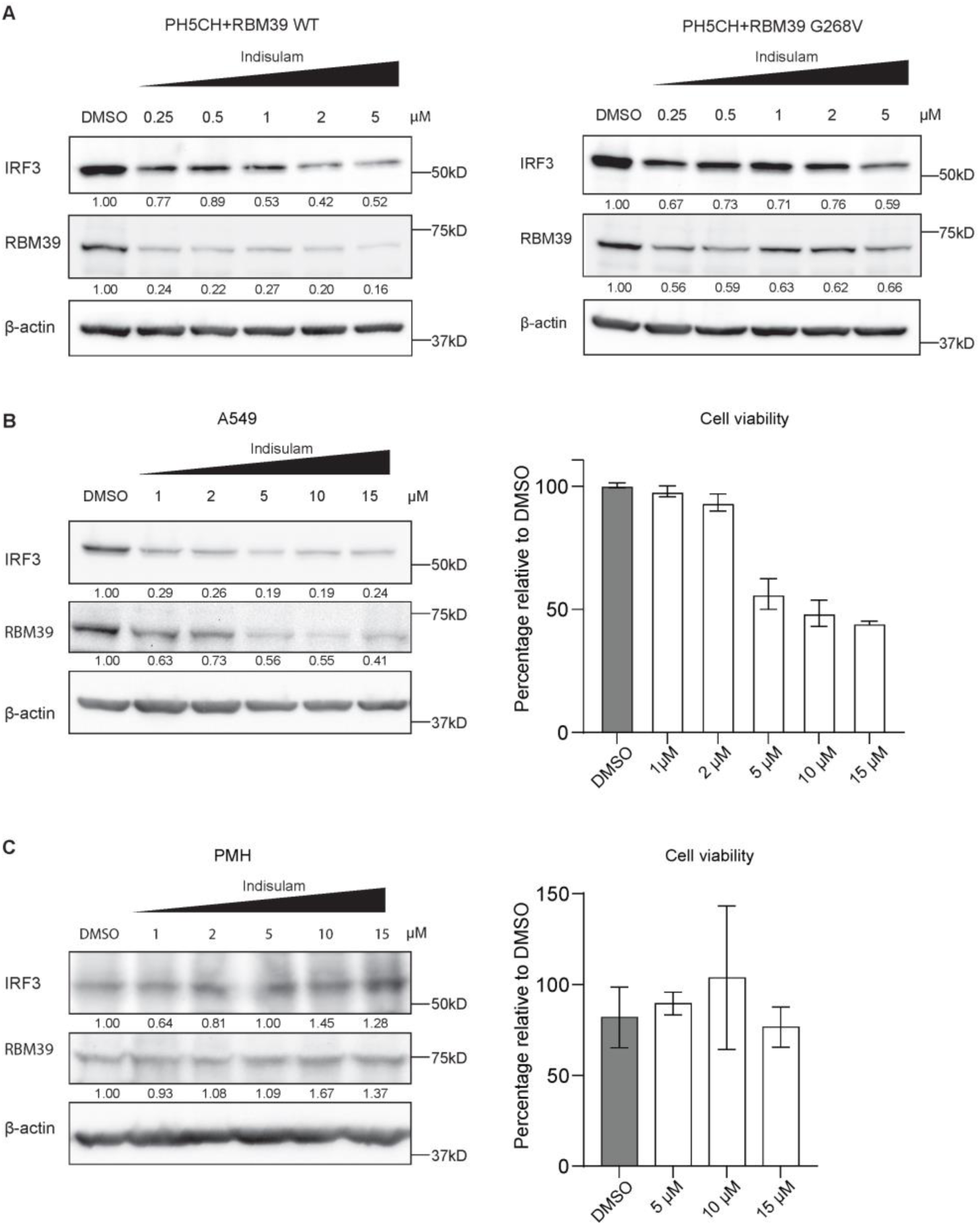
Impact of Indisulam-mediated RBM39 degradation on IRF3 in different cell lines. (**A** to **D**) PH5CH-RBM39 and PH5CH-RBM39 G268V cells (**A**), A549 cells (**B**) and primary mouse hepatocyte (PMH) (**C**) were treated with Indisulam at the indicated concentrations for 48 h. IRF3, RBM39 and β-actin protein expression levels were measured by western blot. Quantification of three independent experiments is shown as average number under the respective bands. (left) Cell viability was measured via CellTiter-Glo luminescent cell viability assay (**B** and **C**, right). Data are from three biological replicates (n = 3), error bars indicate SD.

**Fig. S5.**
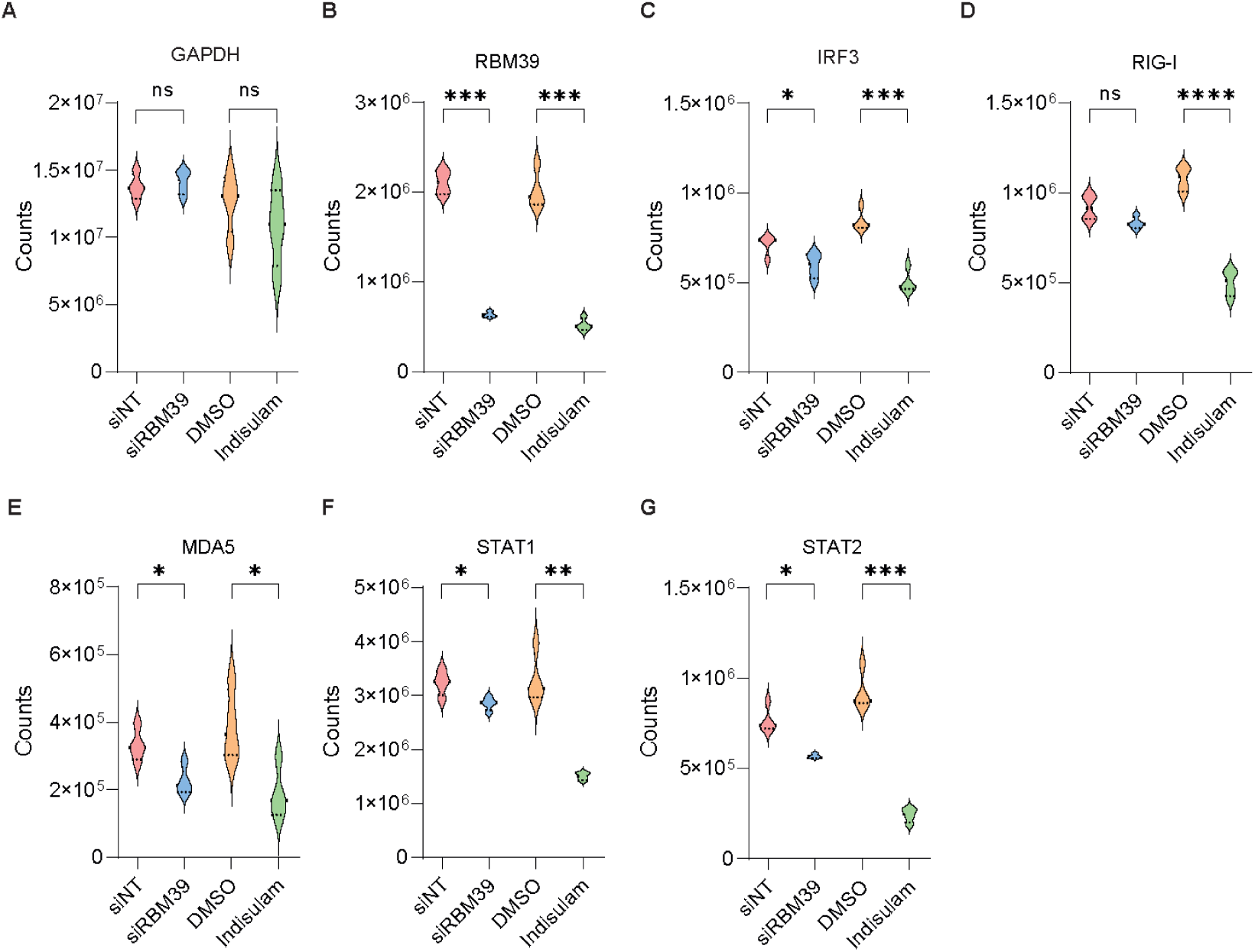
Analysis of individual protein expression levels from proteomics data. The protein level of GAPDH (**A**), RBM39 (**B**), IRF3 (**C**), RIG-I (**D**), MDA5 (**E**), STAT1 (**F**) and STAT2 (**G**) in siRBM39/siNT or Indisulam/DMSO-treated PH5CH samples. Data are derived from three biological replicates (n = 4), error bars refer to SD. Statistical significance was assessed through Welsch’s unpaired *t*-test.

**Fig. S6.**
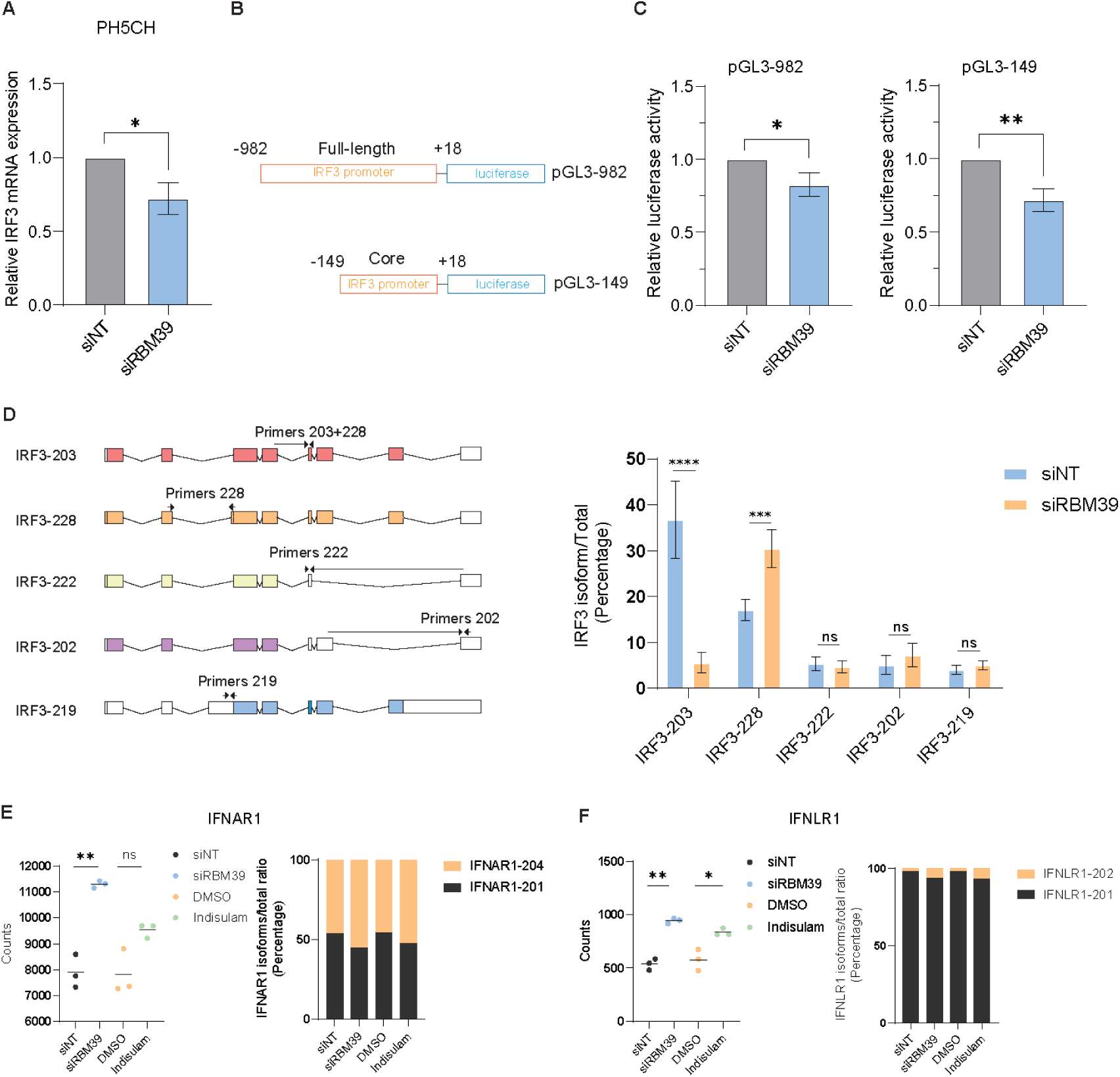
The transcription and splicing of *IRF3*, *IFNAR1* and *IFNLR1 mRNA*. (**A**) PH5CH cells were transfected with siRBM39 or siNT as controls. 48 h after transfection, *IRF3* and *RBM39* mRNA was measured by qPCR. (**B**) Schematic of the firefly luciferase reporter harboring the full-length IRF3 promoter (pGL3-982-firefly) or core IRF3 promoter (pGL3-149-firefly). (**C**) PH5CH cells were transfected with pGL3-982-firefly or pGL3-149-firefly, pGL3-CMV-Gaussia was used as reference. Luciferase activity of firefly was normalized on that of Gaussia. Relative luciferase activity (Relative Light Units, RLU) is shown. (**D**) Schematic of *IRF3* isoforms (left). Specific primers targeting different isoforms are shown as arrows. *IRF3* isoforms/total *IRF3* ratio in PH5CH cells transfected with siRBM39 or siNT as control were identified by qPCR. (**E** and **F**) DEG and DTU analysis of *IFNAR1* (**E**) and *IFNLR1* (**F**) mRNA. DEG and DTU analysis of individual genes were performed using DESeq2 and DRIMseq, respectively. Data shown are from three biological replicates (n = 3), error bars indicate SD. Statistical significance was assessed through Welsch’s unpaired *t*-test.

**Table S5:**
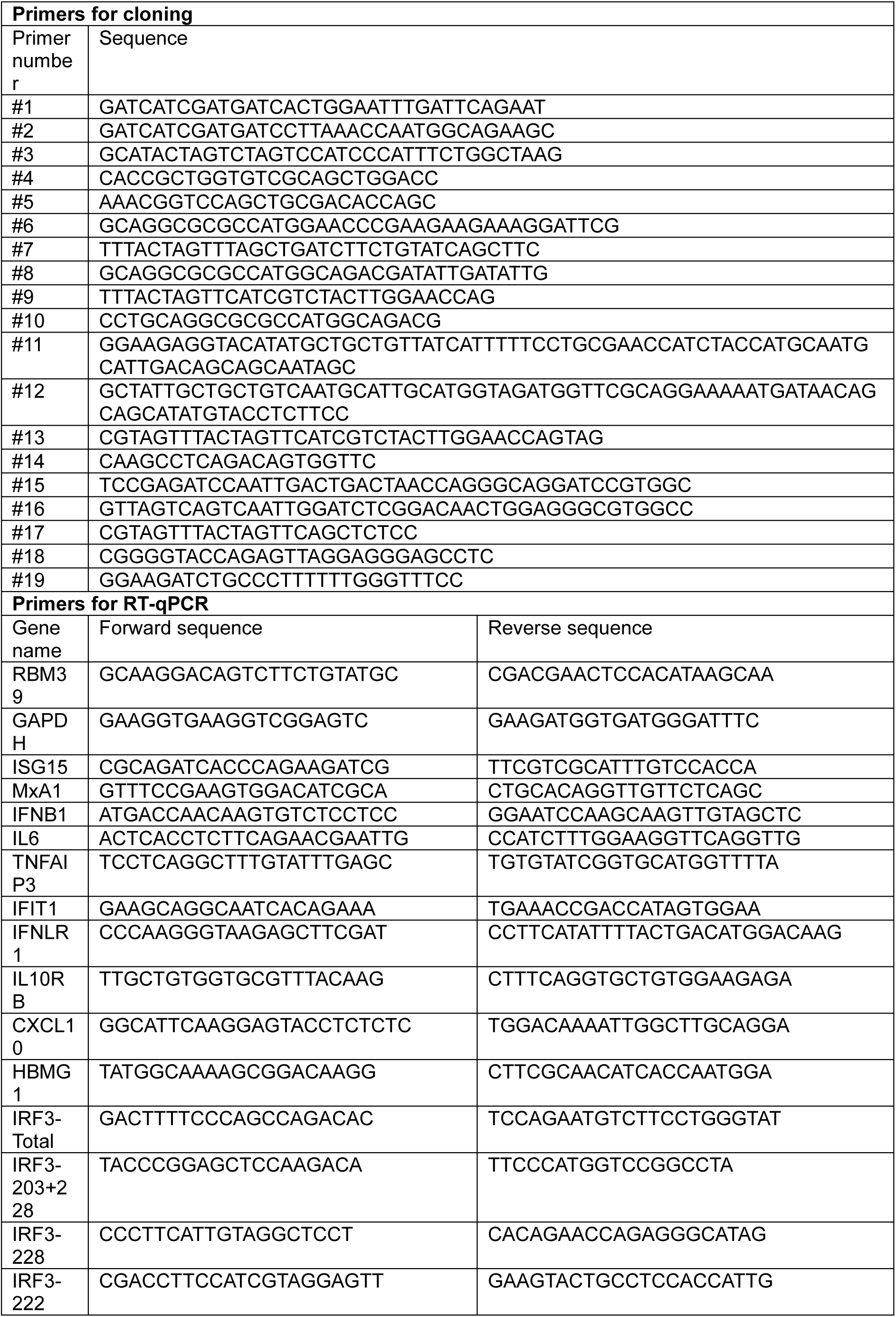
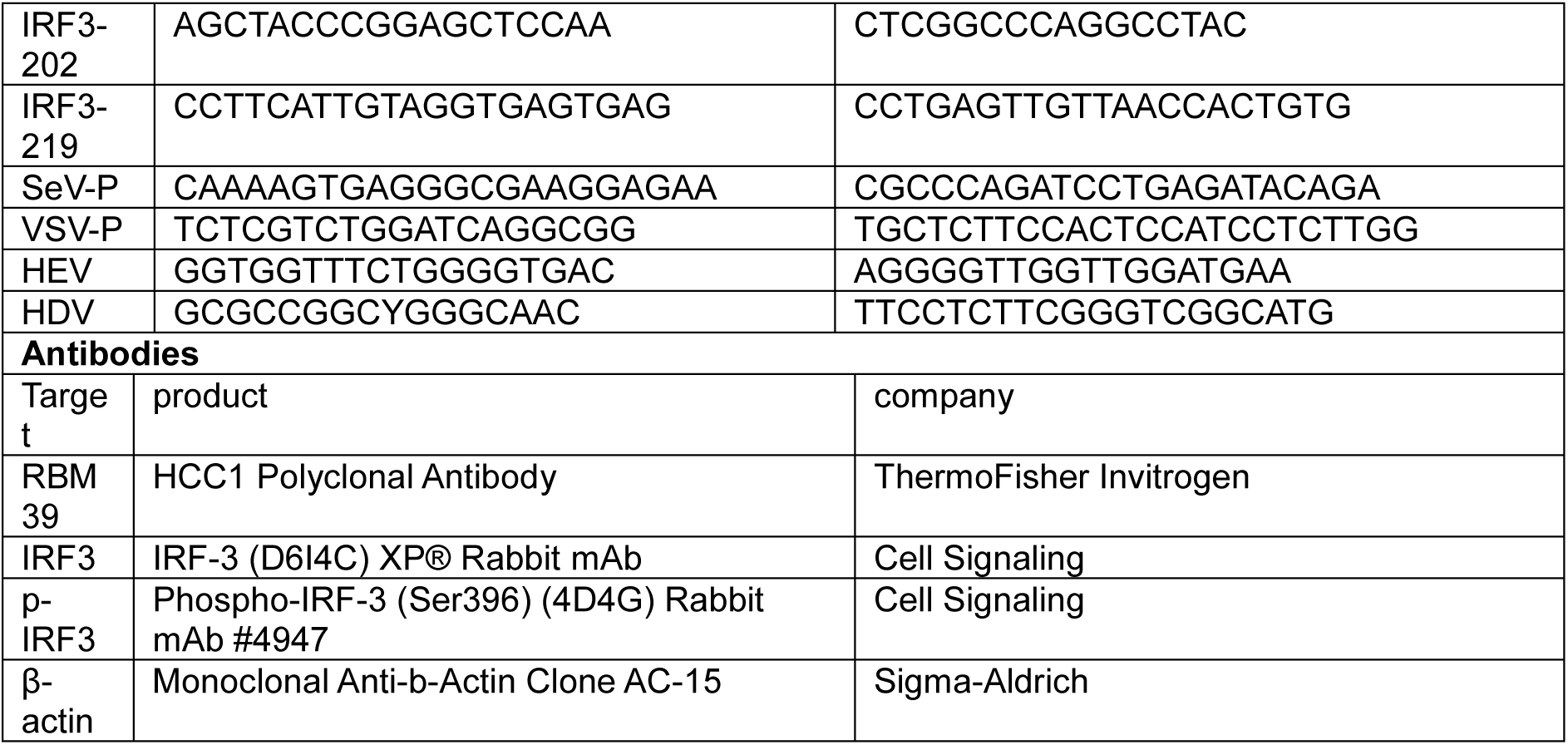
Primers and antibodies.

## Notes

### Competing Interest Statement

The authors have declared no competing interest.

### Summary of Updates

We have repeated the proteomics analysis to achieve higher sensitivity and added some functional analyses to clarify the contribution of factors beyond IRF3 contributing to the phenotype. We finally restructured of the manuscript to include the new data.

