## Supplemental Table 5 for "RBM39 shapes innate immunity through transcriptional and splicing control of key factors of the interferon response"

Primers for cloning

| Primer number | Sequence |
| --- | --- |
| #1 | GATCATCGATGATCACTGGAATTTGATTCAGAAT |
| #2 | GATCATCGATGATCCTTAAACCAATGGCAGAAGC |
| #3 | GCATACTAGTCTAGTCCATCCCATTTCTGGCTAAG |
| #4 | CACCGCTGGTGTCGCAGCTGGACC |
| #5 | AAACGGTCCAGCTGCGACACCAGC |
| #6 | GCAGGCGCGCCATGGAACCCGAAGAAGAAAGGATTCG |
| #7 | TTTACTAGTTTAGCTGATCTTCTGTATCAGCTTC |
| #8 | GCAGGCGCGCCATGGCAGACGATATTGATATTG |
| #9 | TTTACTAGTTCATCGTCTACTTGGAACCAG |
| #10 | CCTGCAGGCGCGCCATGGCAGACG |
| #11 | GGAAGAGGTACATATGCTGCTGTTATCAT  TTTTCCTGCGAACCATCTACCATGCAATGC  ATTGACAGCAGCAATAGC |
| #12 | GCTATTGCTGCTGTCAATGCATTGCATGGTA  GATGGTTCGCAGGAAAAATGATAACAGCAG  CATATGTACCTCTTCC |
| #13 | CGTAGTTTACTAGTTCATCGTCTACTTGGA  ACCAGTAG |
| #14 | CAAGCCTCAGACAGTGGTTC |
| #15 | TCCGAGATCCAATTGACTGACTAACCAGGGCAGGATCCGTGGC |
| #16 | GTTAGTCAGTCAATTGGATCTCGGACAACTGGAGGGCGTGGCC |
| #17 | CGTAGTTTACTAGTTCAGCTCTCC |
| #18 | CGGGGTACCAGAGTTAGGAGGGAGCCTC |
| #19 | GGAAGATCTGCCCTTTTTTGGGTTTCC |

Primers for qPCR

| Gene name | Forward sequence | Reverse sequence |
| --- | --- | --- |
| RBM39 | GCAAGGACAGTCTTCTGTATGC | CGACGAACTCCACATAAGCAA |
| GAPDH | GAAGGTGAAGGTCGGAGTC | GAAGATGGTGATGGGATTTC |
| ISG15 | CGCAGATCACCCAGAAGATCG | TTCGTCGCATTTGTCCACCA |
| MxA1 | GTTTCCGAAGTGGACATCGCA | CTGCACAGGTTGTTCTCAGC |
| IFNB1 | ATGACCAACAAGTGTCTCCTCC | GGAATCCAAGCAAGTTGTAGCTC |
| IL6 | ACTCACCTCTTCAGAACGAATTG | CCATCTTTGGAAGGTTCAGGTTG |
| TNFAIP3 | TCCTCAGGCTTTGTATTTGAGC | TGTGTATCGGTGCATGGTTTTA |
| IFIT1 | GAAGCAGGCAATCACAGAAA | TGAAACCGACCATAGTGGAA |
| IFNLR1 | CCCAAGGGTAAGAGCTTCGAT | CCTTCATATTTTACTGACATGGACAAG |
| IL10RB | TTGCTGTGGTGCGTTTACAAG | CTTTCAGGTGCTGTGGAAGAGA |
| CXCL10 | GGCATTCAAGGAGTACCTCTCTC | TGGACAAAATTGGCTTGCAGGA |
| HBMG1 | TATGGCAAAAGCGGACAAGG | CTTCGCAACATCACCAATGGA |
| IRF3-Total | GACTTTTCCCAGCCAGACAC | TCCAGAATGTCTTCCTGGGTAT |
| IRF3-203+228 | TACCCGGAGCTCCAAGACA | TTCCCATGGTCCGGCCTA |
| IRF3-228 | CCCTTCATTGTAGGCTCCT | CACAGAACCAGAGGGCATAG |
| IRF3-222 | CGACCTTCCATCGTAGGAGTT | GAAGTACTGCCTCCACCATTG |
| IRF3-202 | AGCTACCCGGAGCTCCAA | CTCGGCCCAGGCCTAC |
| IRF3-219 | CCTTCATTGTAGGTGAGTGAG | CCTGAGTTGTTAACCACTGTG |
| SeV-P | CAAAAGTGAGGGCGAAGGAGAA | CGCCCAGATCCTGAGATACAGA |
| VSV-P | TCTCGTCTGGATCAGGCGG | TGCTCTTCCACTCCATCCTCTTGG |
| HEV | GGTGGTTTCTGGGGTGAC | AGGGGTTGGTTGGATGAA |
| HDV | GCGCCGGCYGGGCAAC | TTCCTCTTCGGGTCGGCATG |

Antibody

| Target | product | company |
| --- | --- | --- |
| RBM39 | HCC1 Polyclonal Antibody | ThermoFisher Invitrogen |
| IRF3 | IRF-3 (D6I4C) XP® Rabbit mAb | Cell Signaling |
| p-IRF3 | Phospho-IRF-3 (Ser396) (4D4G) Rabbit mAb #4947 | Cell Signaling |
| β-actin | Monoclonal Anti-b-Actin Clone AC-15 | Sigma-Aldrich |
